# Deep profiling of protease substrate specificity enabled by dual random and scanned human proteome substrate phage libraries

**DOI:** 10.1101/2020.05.09.086264

**Authors:** Jie Zhou, Shantao Li, Kevin K. Leung, Brian O’Donovan, James Y. Zou, Joseph L. DeRisi, James A. Wells

## Abstract

Proteolysis is a major post-translational regulator of biology both inside and outside of cells. Broad identification of optimal cleavage sites and natural substrates of proteases is critical for drug discovery and to understand protease biology. Here we present a method that employs two genetically encoded substrate phage display libraries coupled with next generation sequencing (SPD-NGS) that allows up to 10,000-fold deeper sequence coverage of the typical 6 to 8 residue protease cleavage sites compared to state-of-the-art synthetic peptide libraries or proteomics. We applied SPD-NGS to two classes of proteases, the intracellular caspases 2, 3, 6, 7 and 8, and the ectodomains of the membrane sheddases, ADAMs 10 and 17. The first library (Lib 10AA) was used to determine substrate cleavage motifs. Lib 10AA contains a highly diverse randomized 10-mer substrate peptide sequences (10^9^ unique members) that was displayed monovalently on filamentous phage and bound to magnetic beads via an N-terminal biotin. The protease was allowed to cleave the SPD beads, and the released phage subjected to up to three total rounds of positive selection followed by next generation sequencing (NGS). This allowed us to identify from 10^4^ to 10^5^ unique cleavage sites over a 1000-fold dynamic range of NGS counts (ranging from 3-4000), and produced consensus and optimal cleavage motifs based positional sequencing scoring matrices that closely matched synthetic peptide data. A second SPD-NGS library (Lib hP) was constructed that allowed us to identify candidate human proteome sequences. Lib hP displayed virtually the entire human proteome tiled in contiguous 49AA sequences with 25AA overlaps (nearly 1 million members). After three rounds of positive selection we identified up to 10^4^ natural linear cut sites depending on the protease and captured most of the examples previously identified by proteomics (ranging from 30 to 1500) and predicted 10 to 100-fold more. Structural bioinformatics was used to facilitate the identification of candidate natural protein substrates. SPD-NGS is rapid, reproducible, simple to perform and analyze, inexpensive, renewable, with unprecedented depth of coverage for substrate sequences. SPD-NGS is an important tool for protease biologists interested protease specificity for specific assays and inhibitors and to facilitate identification of natural protein substrates.

## Introduction

Proteolysis is one of the most common post translational modifications (PTMs) and plays essential roles in diverse aspects of cellular functions from protein degradation to specific protein activation^1, 2^. The roughly 600 human proteases—around 2% of the genome—work together to maintain the normal functions and homeostasis of cells and tissues in the body. Aberrant protease activities propagate cancer,^3^ inflammation^4^ and infectious diseases.^5^ Understanding substrate specificities of proteases and their substrates helps define protease functions in cellular processes and provides insights into inhibitor design for both research and therapeutic purposes.

In the past two decades, synthetic peptide libraries, such as Positional Scanning - Substrate Combinatorial Library (PS-SCL)^6^ and Substrate Activity Screening (SAS) methods,^7^ have been used to characterize the linear recognition sequence specificities for proteases. While these are very useful for evaluating 100’s to 1000’s of possible synthetic substrates, they do not deeply sample the possible sequences over the 6 to 8-residue stretch that proteases typically recognize, nor do these random sequences cover exact human sequences. In the past decade, mass spectrometry methods have been developed for identifying intact human protein substrates.^8–12^ While proteomics approaches have enabled a broader understanding of protease substrates on intact proteins, they require significant amounts of lysate, miss low abundance proteins, and miss those simply not expressed in cell lines tested that typically express only half their genomes.^13^

To potentially screen larger and more diverse sequence space, investigators have developed genetically encoded substrate phage^14, 15^ or yeast display libraries.^16, 17^ Degenerate DNA sequences (up to 10^7^) encoding random peptides were fused to a phage or yeast coat protein gene for a catch-and-release strategy or with the assistance of cell sorting, respectively. In the case of substrate phage, the library is bound to an affinity support, exposed to a protease of interest to release, propagated, and enriched for sensitive and resistant clones that are individually sequenced. However, it is difficult to determine the exact proteolytic site by gene sequencing and until now only short 5-6 residue random linear peptide libraries (up to 10^7^ members) have been individually screened, not ones specifically covering the human proteome.

To allow deep substrate profiling we present a next generation of genetically encoded phage display libraries containing either random 10-mers (up to 10^9^ sequence diversity) or human proteome-wide tiled sequences (up to 10^6^ members). We validate these libraries and method on members of the caspase and ADAMs family proteases. Coupling substrate phage display with next-generation sequencing (SPD-NGS) allowed profiling of protease specificity at 10^3^-10^4^-fold greater depth than classical synthetic peptide libraries and identified specific human sequences capable of being cleaved much beyond what has been reported by proteomics. We deployed state-of-the-art positional scoring matrix (PSSM) methods to precisely identify cleavage sites within each selected clone and produce high confidence consensus sequence motifs for the proteases validated by literature. Structural bioinformatics was used to triage the candidate linear substrates for the identification of potential natural protein substrates. We believe these genetically encoded libraries, screening and computational methods provide a simpler, less expensive, and more comprehensive companion to synthetic peptide libraries and proteomics.

## Results

### Substrate phage library strategy, design, assembly and quality control

The overall strategy for SPD-NGS is diagramed in **Figure 1a**. Peptides are displayed as a fusion protein on the surface of M13 bacteriophage. We utilized a monovalent phage display system to avoid avidity and ensure that a single cleavage event per phage is sufficient to release the avidin bound phage. We constructed two <u>s</u>ubstrate <u>p</u>hage <u>d</u>isplay (SPD) libraries, Lib 10AA and Lib hP (<u>h</u>uman <u>p</u>roteome), for complementary and mutually reinforcing purposes (**Figure 1b, 1c, Supplemental Figure 1, 2, 3**). The two libraries were produced using synthetic DNA, and the quality and diversity validated by NGS (**Supplemental Figure 1, 2**). Lib 10AA contains a highly diverse (∼10^9^ unique sequences) and fully randomized ten amino acid substrate segments (**Figure 1b, Supplemental Figure 1**, **3**). The fact that the codon frequency at each position and throughout the 10AA window was uniform and matched the expected input synthetic DNA suggests that there is little cloning or expression bias for the displayed peptides. The strength of the Lib 10AA is to identify highly preferred substrates based on NGS counts from the protease sensitive pool of substrate phage. Consensus cleavage motifs are generated through multiple sequence alignments (MSA) (**Figure 1d**), based on which a positional specific scoring matrix (PSSM) (**Figure 1e**) is derived for high confidence prediction of precise cleavage sites within each selected sequence.

**Figure 1.**
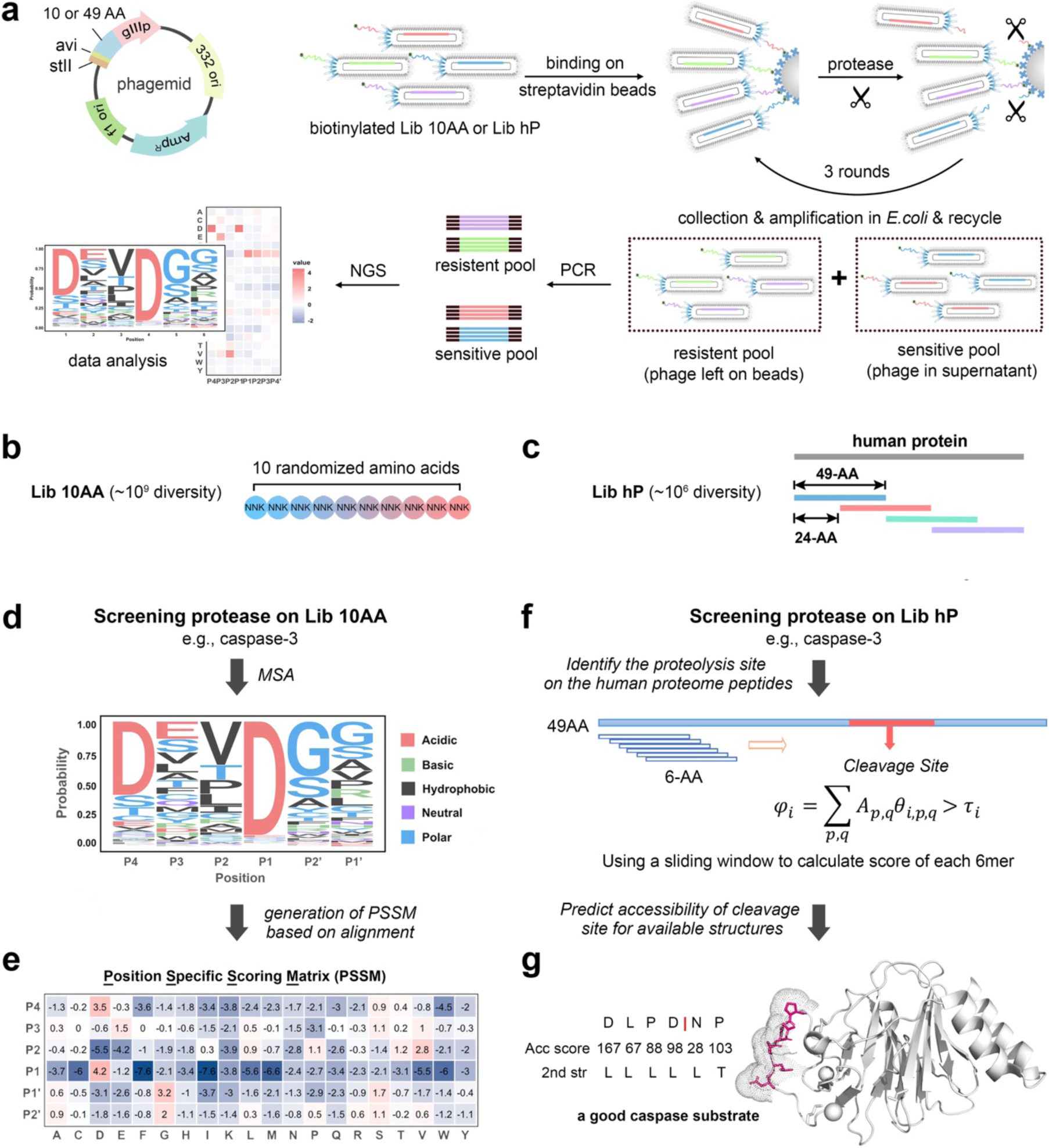
Two-pronged strategy used to identify protease substrates using substrate phage display coupled with NGS (SPD-NGS). (**a**) The schematic illustration of a high-throughput platform for SPD-NGS to profile the protease substrate specificity. (**b**) Lib 10AA contains a highly diverse (∼10^9^ unique sequences) and fully randomized 10AA substrate segment encoded by NNK degenerate codons. N encodes A, T, C, G while K encodes T and G, thus encoding 32 different trinucleotides encompassing all 20 amino acids. (**c**) Lib hP displays tiled peptides covering the human proteome in 49AA blocks. Each protein was computationally divided into 49-AA peptides and overlapped in a 24-AA sliding window to ensure two-fold gap coverage. The resulting 731,724 sequences were synthesized and cloned into a M13 bacteriophage. (**d-g**) Workflow to determine the substrate specificity of a protease in vitro such as caspase-3. A fully randomized 10AA peptide phage library (Lib 10AA) is screened to identify optimal linear peptide sequences from which a scoring function can be generated (**d, e**). A 49AA human proteome tiled library (Lib hP) is screened to identify specific human sequences and sites that can be cut (**f, g**). (**d**) shows a representative 6-residue sequence logo based on SPD-NGS from Lib 10AA for a protease of interest such as caspase-3. The alignment of top 20,000 peptides in this list affords exhaustive generation of a sequence consensus using a R package, ggseqlogo. The data allows us to calculate the probability of each amino acid at each of six positions (P4-P2’). (**e**) The position specific scoring matrix (PSSM) of caspase-3 substrates is generated according to the alignment. (**f**) Determining the precise cut site(s) within each of the positively selected 49AA clones based only on a 6AA segments can be informed by the scoring matrix derived from the Lib 10AA. φ_i_ is a preference score, A is an indicator of peptide sequence (A_p,q_ = 1 if the amino acid at position p of the peptide is q and A_p,q_ = 0 otherwise), τ_i_ is a scoring threshold, specific to each protease and θ_i,p,q_ is positive if a certain amino acid is preferred and otherwise negative.^24^ (**g**) Solvent accessibility (Acc score) and secondary structure (2^nd^ str) of the potential cleavage site, calculated by DSSP (Define Secondary Structure of Proteins) program, are used to enhance the prediction of whether the folded protein is a potential protease substrate (adapted from PDB: 4N7I).

The Lib hP library contains nearly complete coverage of human proteome sequences in 49AA blocks, tiled with 25AA overlaps for duplicate gap coverage containing ∼730,000 individual sequences (**Figure 1c**, **Supplemental Figure 2, 3**). The Lib hP has the advantage that all substrate sequences are from the human proteome and it provides direct information about what linear sequences can be cut in the human proteome. The Lib hP also provides broader coverage than one can typically achieve by proteomics or RNAseq because no single cell line expresses the entire genome (**Supplemental Figure 2e, 2f**). When coupled with sequence and structural bioinformatics the Lib hP data identifies candidate substrates for more detailed analysis at the protein level (**Figure 1g**).

Each substrate phage displays an avi-Tag on in its N-terminus to permit quantitative biotinylation and immobilization on streptavidin magnetic beads (**Figure 1a**). For the input library we biotinylated the displayed peptide *in vitro*. We confirmed the quantitative presence of biotin attached to the peptide displayed on phage using a phage ELISA (**Supplemental Figure 4a, 4b, 4c**). In subsequent rounds of selection to facilitate biotinylation, we passaged enriched substrate phage pools into *E. coli* XL-1 Blue cells that we engineered to express intracellular biotin ligase BirA with pBirAcm, an engineered pACYC184 plasmid with an IPTG inducible birA gene, so biotinylation can be simultaneously done during phage amplification **(Supplemental Figure 4c, 4d**).

### Using SPD-NGS to profile the linear specificity of caspases

Caspases are of high biological interest because of their powerful roles in cell death and differentiation.^18^ These are excellent proteases to validate the SPD-NGS approach because they have been extensively studied using traditional synthetic peptide libraries and proteomics. We began by profiling caspase-3 (**Figure 2**). It is challenging to know *a priori* the ideal concentration of enzyme, incubation time, and number of rounds of selection for optimal profiling. Thus, we determined these empirically by covering a 1000-fold range of enzyme concentrations and monitoring enrichment as a function of multiple rounds of selection. Briefly, avidin magnetic beads were incubated with the Lib 10AA phage, washed extensively to remove non-bound phage and freshly expressed caspase-3 was added for 30 min at room temperature at enzyme concentrations varying from 1-1000nM. We measured the number of released phage by counting infectious units as a function of round of selection (**Supplementary Figure 5**). The number of released phage over the untreated substrate phage beads increased by 10-1000 fold as a function of round of selection and enzyme concentration suggesting strong positive selection.

**Figure 2.**
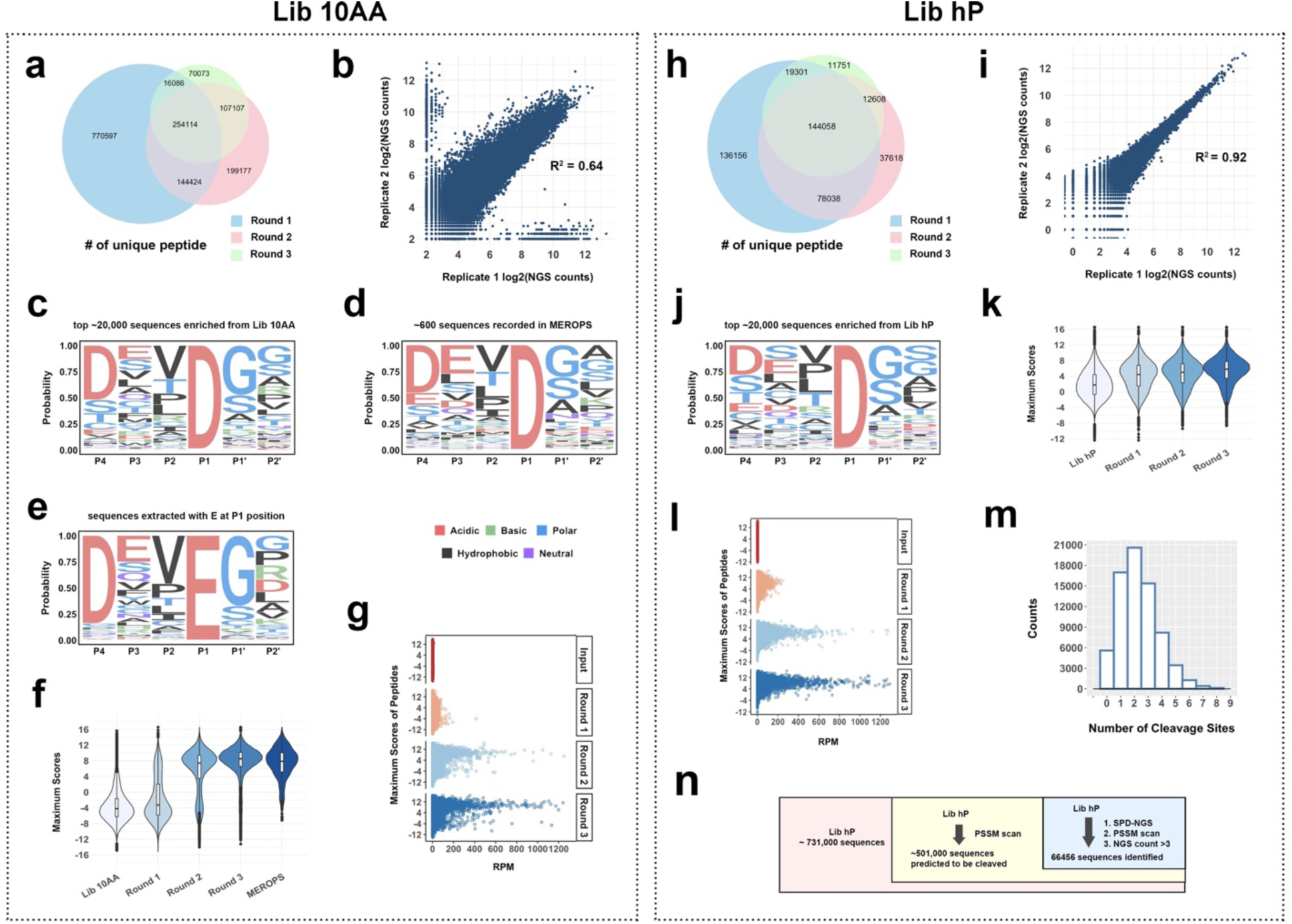
Validation of SPD-NGS libraries to identify protease specificity for caspase-3. (**a**) The Venn diagram of unique peptides identified from Lib 10AA as a function of rounds of selection. The numbers decrease due to increasing enrichment of better substrates with rounds of the selection. (**b**) The relatively strong correlation between two biological replicates (R^2^=0.64) indicates the reproducibility of the selection strategy. (**c**) The Sequence Logo for caspase-3 substrates generated by aligning the top 20,000 sequences based on counts identified from screening Lib 10AA (Round 3). (**d**) The Sequence Logo for caspase-3 substrates generated from substrates compiled in MEROPS database. (**e**) A similar motif is obtained by selecting only proteolytic sites with a glutamate (E) at P1 position as seen previously by N-terminomics data of caspase-3.^23^ (**f**) Violin plot for distribution of scores of 10AA sequences for the input library, Round 1, 2, 3 outputs, and the scores of peptides in MEROPS proteomics database. The peptides with higher scores get progressively enriched in Round 2&3 and Round 3 match scores seen in MEROPS. (**g**) The plot of maximum score vs RPM for peptides in input library and Round 1, 2, 3 outputs. The peptides with higher scores get enriched faster. (**h**) The Venn diagram of unique peptides identified from Lib hP decreases with each of three rounds of the selection as enrichment increases as was seen for Lib 10AA. (**i**) The correlation for two biological replicate Round 3 for the Lib hP. A stronger correlation (R^2^=0.9) between two biological replicates indicates the reliability of the selection strategy and the reduced starting library size for Lib hP (∼10^6^) compared to Lib 10aa (∼10^9^). (**j**) The Sequence Logo of caspase-3 substrate consensus generated by top 20,000 cleavage events identified from Lib hP. (**k**) Violin plot of the maximum scores of the Lib hP input library, and progressive enrichment of substrates as one progresses from output of Round 1, 2, and 3 which then aligns with MEROPS seen in 3d. (**l**) The plot of maximum score vs NGS RPM for peptides in input library and Round 1, 2, and 3 outputs. The peptides with higher scores get enriched faster. (**m**) Distribution of frequency observed as a function of number of cuts in the 49AA peptides. Most have one cut but some have multiple that decays monotonically. (**n**) Venn diagram of the library (Lib hP), the peptides passing PSSM scan and the peptides identified from SPD-NGS.

After each round of selection, NGS was applied to sequence the protease sensitive pools. In a pooled substrate screen such as this, substrates should enrich based on their relative catalytic efficiencies for hydrolysis (k_cat_/Km). Thus, with increasing rounds of selection we would expect that the number of unique reads to decrease and NGS counts per sequence to increase as good substrates are selected over poor ones. Indeed, from an average of 500,000 NGS reads per round we identified roughly 377k, 272k and 187k unique sequence reads from Rounds 1, 2, and 3, respectively (**Figure 2a**). The selection for better substrates was further supported by the Venn diagram showing that generally more than 60% of the substrates identified in a later round were also present in the set from the previous round; virtually all ∼350k substrates found in Round 3 were also found in the set of substrates found in Round 2 and similarly for Round 2 contained in Round 1. Two biological replicates were performed for each of the three rounds of positive selections for each of the three enzyme concentrations; these showed a strong correlation (R^2^=0.64) indicative of the good reproducibility of the selections (**Figure 2b**). This is not a perfect correlation which is likely the result of NGS sampling only a small portion (∼10^5^-10^6^) of the possible unique sequences due to high diversity of the starting Lib 10AA (∼10^9^). The reads per million (RPM) for each unique sequence ranged from a few to over 1000, indicative of the wide dynamic range and variability in rates of hydrolysis for individual substrates.

We next analyzed the enrichment based on Z-scores over the starting library for each of the 20 amino acids as a function of round of selection and enzyme concentration used over the 10AA substrate window (**Supplemental Figure 6**). We see obvious enrichments for particular amino acids (Asp, Glu and Gly) and de-enrichment for others (Leu and Arg) especially in the middle of the 10AA substrate linker in Round 3 **(Supplemental Figure 6c, 6f, and 6i**). Also, simple analysis of residue preferences across the 10AA substrate window shows a strong selected amino acid preference in the middle 6-7 residues favoring Asp, Glu, Val, and Gly and disfavoring Phe, His, Lys, Arg, Trp (**Supplementary Figure 6, 7a**). These preferences increase with increasing round of selection and enzyme concentration further suggesting positive selective pressure for some residues and not others. These data are consistent with structural data showing caspase-3 binds a six-residue linear stretch of peptide (**Supplemental Figure 7b**) with known subsite preferences such as P4(acidic)P3(acidic)P2 (hydrophobic)P1 (acidic) and P1’ (small). A simple multiple sequence alignment (MSA) of the top 20,000 unique peptides cleaved by caspase-3 generated Z-score enrichment values over the input library for a six-residue window (**Supplemental Figure 7c**). The top scoring residues were reflected the classic caspase-3 motif, DEVD/GG. We can also represent this data in a familiar sequence Logo that expresses the probability of each amino acid at each of the six positions (**Figure 2c**). This Logo based on ∼20,000 sequences is remarkably close to what has been seen from synthetic peptide libraries and proteomics compiled in the MEROPS database of about ∼600 sequences^19–22^ (**Figure 2d; Supplemental Figure 7d**). It is also known that caspase-3 can cleave with Glu at P1.^23^ Indeed, a sequence logo derived from positively selected peptides containing a Glu at P1 position after MSA (**Figure 2e**) was virtually identical to the motif for Asp (**Figure 2c**) (DEVE/GG versus DEVD/GG).

### Generating an unbiased PSSM for caspase-3 using consensus data from Lib 10AA

To more rigorously compare the quality of substrates and to predict the precise cleavage sites on the 49AA sequences identified from Lib hP, we developed a traditional position specific scoring matrix, PSSM. A peptide is predicted to be cleaved by caspase-3 if:

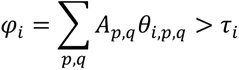

Where φ is a substrate score, A is an indicator of peptide sequence (*A_p,q_* = 1 if the amino acid at position *p* of the peptide is *q* and *A_p,q_*= 0 otherwise), and τ*_i_* is a scoring threshold, specific to the protease (**Figure 1c; Supplemental Figure 8**).^24, 25^ In most cases, τ*_i_* is 0 although one can set a higher threshold for only good substrates. We arbitrarily assumed that the contribution of each peptide position to selective recognition is additive. For caspase-3, our model takes into account a 6AA peptide: positions P4-P2’, according to alignment and caspase-3 crystal structure (**Supplementary Figure 7b**).

The PSSM model, described further in the Materials and Methods, generated a quantitative score for 6mer substrate peptides scanned over the selected sequences. Applying the PSSM to the caspase-3 NGS dataset from each round at the highest concentration (1 µM, 30 min treatment), we generated violin plots showing the distribution of the number of peptides versus the maximum PSSM score a 6AA peptide within the 18AA window (4AA on each side + 10AA) (**Figure 2f**). The majority of sequences in the input library had relatively low scores and there was little change at Round 1. After Round 2 the majority were high scoring sequences, and by Round 3 almost all sequences were of high scores. Remarkably, the Round 3 violin plot had an even slightly higher average PSSM score than the violin plot for caspase-3 cut peptides from the MEROPS database. The violin plots are generated based on more than 4 X 500,000 seuences. If we set τ_i_ for caspase-3 to be a zero threshold, the PSSM model derived from our data produced a true-positive rate >95% of the MEROPS database. Another way to view the enrichment as a function of round of selection was simply by plotting the number of NGS sequence counts (represented by reads per million (RPM)) for each peptide versus their corresponding PSSM score (**Figure 2g**). There is a clear increase in the number of highly scoring peptides as the selection proceeds through the three rounds relative to the unselected input.

### Selection of caspase-3 substrates from Lib hP

We next applied the SPD-NGS protocols that were optimized for caspase-3 on Lib 10AA to the human focused Lib hP. As for Lib 10AA, we conducted three rounds of selection and similarly see the number of unique selected sequences decreases significantly from roughly 370,000 to 28,000 to 20,000 in going from Round 1 to Round 3 (**Figure 2h**). Again, about 80∼90% of the sequences found in the Round 2 pool were present in Round 1 and similarly for Round 3 captured in Round 2 suggesting the better sequences were winning over poorer sequences. We also analyzed the data by filtering out low abundance peptides having fewer than 3 reads and obtained similar results albeit with almost 2-fold lower number of candidate substrates (**Supplemental Figure 9a**). However, the overlap per round was more suggesting the unfiltered set would have more false positives so we retained a threshold of >3 reads.

We also tested the reproducibility of the selections by conducting two biological replicates for three rounds on Lib hP. A plot of the number of NGS counts per identical peptide sequence for Replicate 1 versus 2 (**Figure 2i**) showed a linear and remarkably high correlation (R^2^=0.92). We believe the higher correlation coefficient seen among these replicates from Lib hP is a result of the 1000-fold lower diversity of this library compared to Lib 10AA allowing more complete capture of positive clones.

Subjecting the top selected 49AA sequences to a simple MSA to get a reliable consensus is much more challenging given the many possible alignments one could generate in a 49AA window from Lib hP versus the 10AA window in Lib 10AA. However, it is reasonable to assume the sequence preferences for caspase-3 are the same whether cut in a 10AA window or a 49AA window. Thus, we applied the six residue PSSM derived from Lib 10AA to Lib hP for identifying potential cleavage site(s) and derived a sequence Logo for top 20,000 sequences as we did for Lib 10AA (**Figure 2j**). To do this we calculated the score (φ_i_) of each possible 6mer on every cleaved sequence in the 49AA NGS dataset after 3 rounds of selection and considered it a cleavage site if φ_i_ > 0. Indeed, the sequence Logo derived from these data (**Figure 2j**) is remarkably close to that seen in the consensus from Lib 10AA and MEROPS database (**Figure 2c, d**). Violin plots were generated from these and revealed a consistent increase in the average NGS counts as the selection proceeded from Round 1 to 3 (**Figure 2k**). The change in distribution as a function of rounds is also evident from increase in counts for specific sequences as that selections proceeded (**Figure 2l**) as was seen in the Lib 10AA data (**Figure 2g**) The increases in NGS counts per round for Lib hP was less dramatic than seen for Lib 10AA, and likely reflects the fact that the average number of cuts within the 49AA window was 1.5 and ranged from 0 to six (**Figure 2m, Supplementary Figure 8b**). We also tested if we could simply apply the PSSM score derived from the Lib 10AA to identify potential cleavage sites in the human proteome. Applying the PSSM on the Lib hP sequences identified ∼500,000 peptides that could potentially be cleaved by caspase-3 which was about 10-times more than the actual ∼ 60,000 peptides (NGS counts >3) enriched from SPD-NGS screening of the Lib hP (**Figure 2n**). Although the MSA and PSSM are useful for identifying the most probably cut site within a selected clone they are not a substitute for the experimental selections.

We next compared the unique human sequences identified by SPD-NGS (∼60,000) to those found by mass spectrometry (∼600). As seen in the Venn diagram nearly 64% of the proteomics database is found in the Lib hP data set (**Figure 3a**). There are ∼700,000 Asp in the human proteome and only 4% are cut in the Lib hP indicating much more is required than simply a P1 Asp for efficient cutting. We also find about 100-fold more substrates than have been identified by mass spectrometry. This could be for a number of reasons. Mass spectrometry of lysates would typically miss the ∼4000 membrane and extracellular proteins that would not be cleaved by intracellular caspases. In addition, cells only express about half of their genomes and low abundance proteins would be missed by mass spec methods (**Supplemental Figure 2e, 2f**).

**Figure 3.**
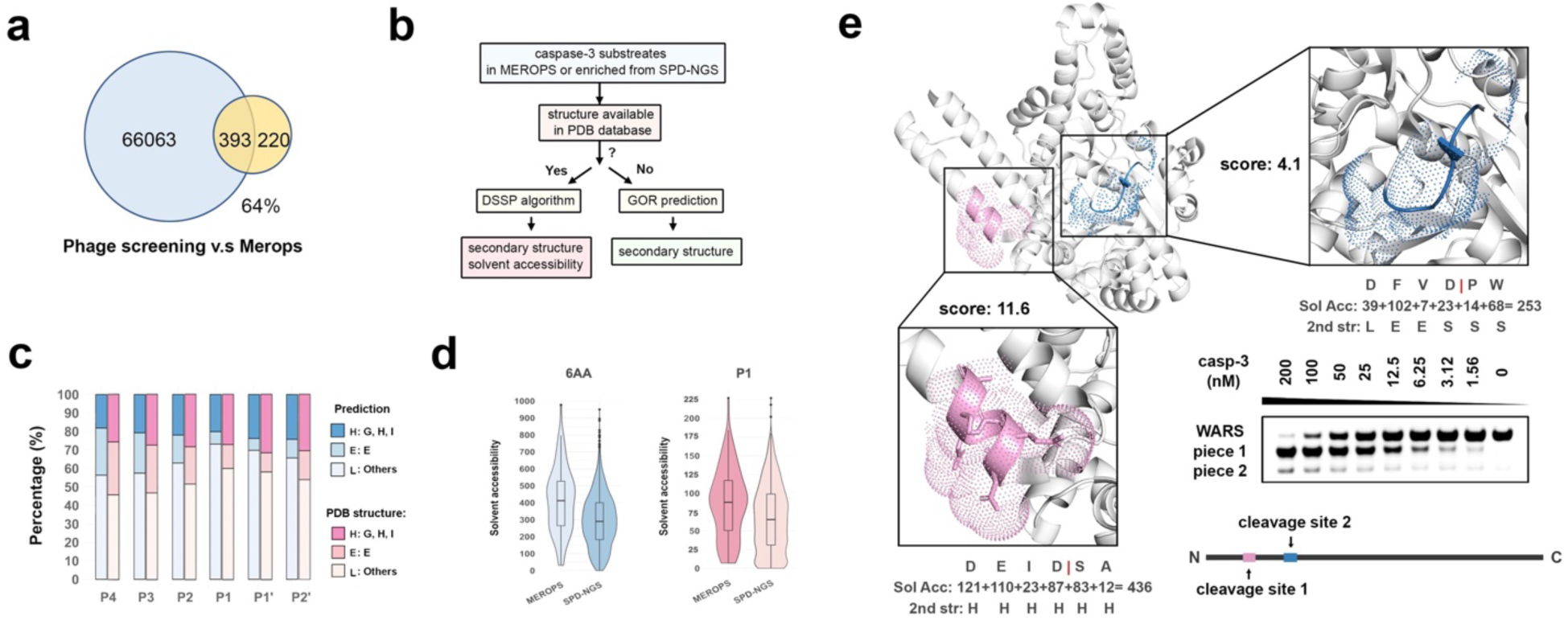
Triaging the candidate linear substrates by structural bioinformatics. (**a**) SPD-NGS identifies 64% of caspase-3 substrate sequences recorded in MEROPS database (only protein substrates are used here). (**b**) The workflow showing to calculate the secondary structure and solvent accessibility of the cleaved peptides enriched in SPD-NGS in a protein context (if PDB structures available) as supplement to determine whether the protein where the peptide is derived from a potential substrate. (**c**) Validation of an unreported protein substrate of caspase-3, tryptophanyl-tRNA synthetase (WARS) identified by SPD-NGS. The ribbon diagram and predicted solvent accessibility of the WARS protein (PDB structure 5UJJ) shows the two sites (1 and 2) and solvent accessibility scores (DSSP) of 60 and 37, respectively. SDS gel shows the most accessible Site 1 is cleaved at lower caspase-3 concentrations compared to less accessible Site 2. (**d**) caspase-3 has a known structural preference for cutting loop>helix>sheets and this matches structural bioinformatics for sites identified from Lib hP. The DSSP (structure available) or GOR (Garnier-Osguthorpe-Robson, structure unavailable) method were used for 2^nd^ structure prediction. (**e**) The violin plots of solvent accessibility for the residue at P1 identified from Lib hP (structure available) and recorded in MEROPS database. The proteolysis recorded in MEROPS database are real cleavages, most of which are exposed.

### Using Lib hP data plus solvent accessibility to predict new protein substrates

Although the Lib hP can broadly identify what can be cut when exposed, we anticipate the majority of these sequences may not be accessible to the protease in the native folded protein substrate. Thus, to help triage the candidate linear substrates, we found it useful to apply a global modeling and surface accessibility score, DSSP (Dictionary of Secondary Structure of Proteins)^26^, to rank cleavage sites based upon surface accessibilities. The application of DSSP requires a 3D structure which only applies to a subset. For those substrates where PDB structures are not available, we estimated their secondary structure using the Garnier-Osguthorpe-Robson (GOR) algorithm^27^ (**Figure 3b**). Previous proteomics studies have shown that caspases have a preference for cleaving loops over helices over sheets in native proteins^19^. When we apply our candidate sequences from the Lib hP to secondary structure prediction algorithms or look at the structures in a protein context, we find a similar preference for loop, followed by helix and then sheet (**Figure 3c; Supplemental Figure 9c**). A plot of the DSSP values for those substrates (6mer) or residues at P1 position of known structure shows a broad gaussian distribution (**Figure 3d**). We made the same calculation for those substrates found in the proteomics dataset for the caspase-3 (**Appendix Table S1)** and find a higher mean score (**Figure 3d**) which emphasizes the role of accessibility.

Realizing that site accessibility is critical for cleavage by caspase-3,^28^ we scanned our dataset to identify potential substrates with a known structure. For caspase-3, we filtered the sequence list based on the intracellular location expected for substrates of caspase-3. For a solved structure or a comparative model, we used the DSSP program to assess secondary structure (mapping results ‘H’, ‘G’ and ‘I’ to helix; ‘B’ and ‘E’ to sheet; and ‘S’, ‘T’ and ‘L’ to loop) and solvent accessibility.^26^ When a structure or model was not available, we used sequence-based algorithms to predict secondary structure (**Appendix Table S2-6**).^27^

We choose to test a protein that had not been reported before, tryptophan-tRNA ligase (WARS). We identified two candidate sites in the WARS protein with solvent accessibility of 436 (DEID|SA) and 253 (DFVD|PW) (**Fig. 3e; Supplementary Figure 10a, b**). The 436 site was located in a helix and had a PSSM score of 12.1, whereas the 253 site was located in a sheet had lower PSSM score of 6.4. After expressing the WARS protein in E coli, we treated it with different concentrations of caspase-3 and used SDS-PAGE to probe for these two cleavage events. Indeed, we observed two cleavage sites and confirmed the molecular weight of each piece with LCMS (**Supplemental Figure 10**). Moreover, the DEID/SA sequence was cleaved much more rapidly than the DFVD/PW site as predicted by both higher solvent accessibility and PSSM score.

### Generalization of SPD-NGS to other caspases

Having built and validated the two complementary substrate phage libraries, and established a work flow to collect and analyze the SPD-NGS data for caspase-3, we expanded the approach to other caspases involved in cell death for which extensive proteomics work has been applied including caspases- 2, -6, -7 and -8. We conducted the two-step substrate phage selections with Lib 10AA and Lib hP as performed on caspase-3, using freshly expressed recombinant active caspases-2, -6, -7, and -8 (**Supplemental Figures 11-15**). To routinely produce the active caspases, it was necessary to remove the pro-domains. The caspase-2 used in this study contained no N-terminal caspase recruitment domain (known as ΔCARD-caspase-2) and caspase-8 had the death-effector domain removed (known as ΔDED-caspase-8). The caspase activities were validated on fluorogenic substrate Ac-DEVD-R110 (**Supplemental Figures 15**). As for caspase-3, each of the selections using Lib 10AA showed increasing enrichments as the three rounds proceeded. This permitted us to generate a PSSM for each caspase, and to construct sequence logos (**Supplemental Figures 11-14**). Similarly, the selections with the Lib hP proceeded with enrichment through the three rounds, and application of the Lib 10AA PSSM allowed generation of a sequence logos from both Lib 10AA and Lib hP data.

These experiments generated a massive amount of data which was compiled to allow comparison among the apoptotic caspases both from the SPD-NGS and existing proteomics data (**Figure 4**). We first compare the Z-score enrichment heatmaps for all five caspases generated from the Lib 10AA data to obtain general comparative features (**Figure 4a; Supplemental Figure 11-14**). Using hierarchical clustering which incorporates all of the data it is apparent that caspases-3 and -7 are most similar; this is not surprising given their central role as executioner caspases. Both have the dominant DEVD|GG motif and are almost indistinguishable in other general subsite details. Caspase-2 is more closely related to these two executioners except for subtle differences at P2 showing a dominant DESD|GG motif. The biggest difference is the preference for polar residues (S, T, R) at P2 compared to the V at P2 for caspases-3 and -7. This raises the possibility that caspase-2 is more executioner like in its function compared to caspases-6 and -8.^29^ In contrast, caspases-6 and -8 are more similar to each other having a dominant VEVD|GG and LETD|GG motif, respectively. These two differ most at P4 from the executioners. Although caspase-6 has long been thought to be more executioner-like because of higher sequence homology, more recent studies suggest it plays a more independent role and may be more of an initiator like caspase-8.^30, 31^ Interestingly the heat maps also show that although Asp is clearly dominant for all the caspases at P1, the next most enriched residue is Glu suggesting it should not be ignored at possible cleavage sites assuming other subsites are strongly satisfied.

**Figure 4.**
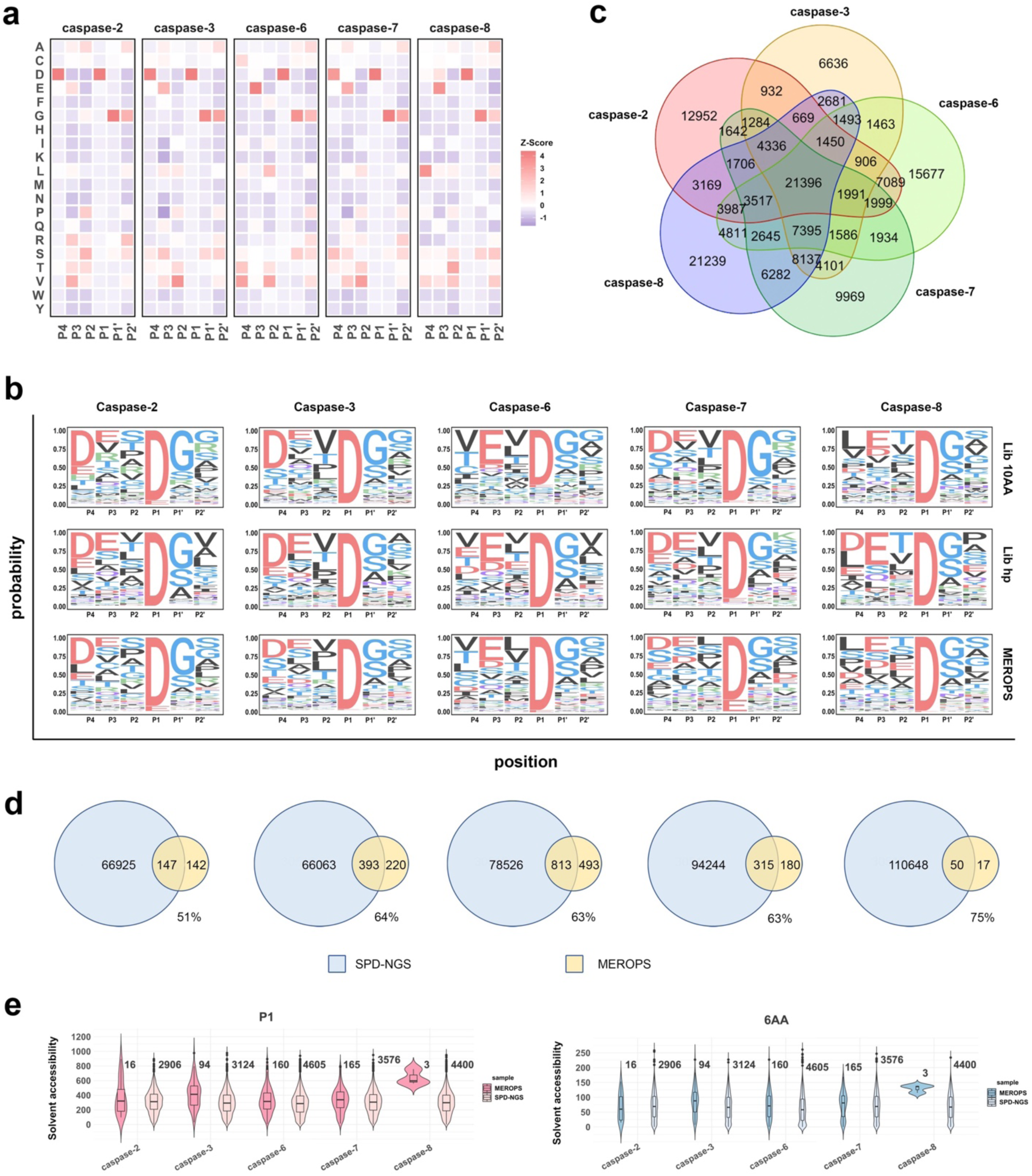
SPD-NGS for profiling substrates of other caspases. (**a**) Z-score Heatmap showing positional enrichment or de-enrichment of each amino acid at P4-P2’ compared to the input libraries (Lib 10AA). (**b**) Sequence Logos of substrates for purified caspases -2, -3, -6, -7, and -8 generated from the top 20,000 sequences selected by SPD-NGS from Lib 10AA, Lib hP, and for 300-1300 sequences reported in the MEROPS database. (**c**) Venn diagrams of the sequences selected from Lib hP (ranging from 43k to 127k) and those reported in MEROPS database (ranging from 300 to 1300). The SPD-NGS from the Lib hP captured 30-60% of those reported in MEROPS and generally identified 100x more cut sites. (**d**) Venn diagrams for the overlap of substrates found by SPD-NGS and proteomics of individual caspases. (**e**) Venn diagrams of the sequences identified to be substrates of different caspases. Note that only 5-10% are common to all, and 20-30% are unique to each.

We generated three sequence logos for each of the caspases from the Lib 10AA, Lib hP and MEROPS database for comparison (**Figure 4b**). There is remarkable agreement in the patterns seen when comparing the data obtained by either SPD-NGS method to proteomics for each caspase. When we compare across caspases we come to the same interpretation we did from the heatmap data in **Figure 4a**; caspases-2, -3 and -7 group together as do caspases-6 and -8. The biggest distinction between these two groups is that caspases-2, -3, and -7 prefer D at P4 over E at P3, and caspases-2 and -8 prefer a hydrophobic (V or L) at P4 and stronger preference for E at P3.

We next analyzed specific human substrates found in in the SPD-NGS Lib hP dataset for all five caspases-2, -3, -6, -7 and -8 uploaded into **Appendix Table S2-6**, respectively. These resource tables present gene name of each substrate, the 49AA sequence, cleavage site predicted, PSSM score, subcellular location, PDB structures if available, solvent accessibility, and secondary structure of the predicted cleavage site. For caspases-2, -3, -6, -7, and -8 we identified ∼60K unique sequences having >3NGS reads from ∼10K proteins, respectively. The two to three-fold range in numbers of unique substrates identified is narrow and generally tracks with the relative specific activity of each caspase. We also compared the overlap of substrates identified in Lib hP among the five caspases (**Figure 4c**). Interestingly there was an overlap of ∼20,000 common substrates suggesting some redundancy, but a larger set of 6,000-20,000 that were unique to each caspase. We do not believe the unique sets are due to sampling issues since we found high reproducibility in the biological replicates (**Figure 2i**).

In **Figure 4d**, we compare the overlap of substrates found by SPD-NGS from the Lib hP and corresponding proteomics data sets. Remarkably, on average more than 50% of the substrates identified by proteomics are contained in the respective SPD-NGS datasets. The number of human substrates identified from the caspase-3 Lib hP dataset was 66,000 substrates compared to about 600 so far identified by proteomics, a difference of 400-times more in the SPD-NGS dataset. We believe greater accessibility of the linear peptides in the Lib hP versus accessibility in the folded proteins in the proteome is the major factor that accounts for much of this difference. To compare these datasets based on accessibility we filtered the SPD-NGS dataset for those substrates where a structure is available in the Protein Data Bank to allow calculation of accessibility by DSSP (values shown in **Appendix Table S2-6**). A plot of the DSSP value for those substrates of known structure shows a broad gaussian distribution (**Figure 4d**). We made the same calculation for those substrates found in the proteomics dataset for the five caspases (**Appendix Table S1)** and find a higher mean score (**Figure 4e**). We would expect that the true protein substrates would have a distribution more like the proteomics data. Since the sample sizes of true protein substrates for caspases are small, finding an accessibility threshold for the linear peptides in their native folded protein is rather challenging. Substrates identified in SPD-NGS from the Lib hP with a P1 accessibility > 25 and 6AA stretch accessibility > 100 would be top candidates for further detailed validation in vitro.

### Generalizing the protocol to the ADAMs family sheddases — ADAM10 and ADAM17

ADAM10/17 are membrane bound proteases involved in shedding of extracellular domains in signaling (**Figure 5a**). They are produced as inactive zymogens that become activated. They are more highly activated in the tumor microenvironment^32^ and identifying substrates is of high interest in cancer. Although only 30 substrates are known for ADAM17, many are of high biological interest such as TNF, NOTCH1, APP, EGF. The specificity of ADAM 10 and 17 has been studied using a library of 200 synthetic 10-mer peptides.^33^

**Figure 5.**
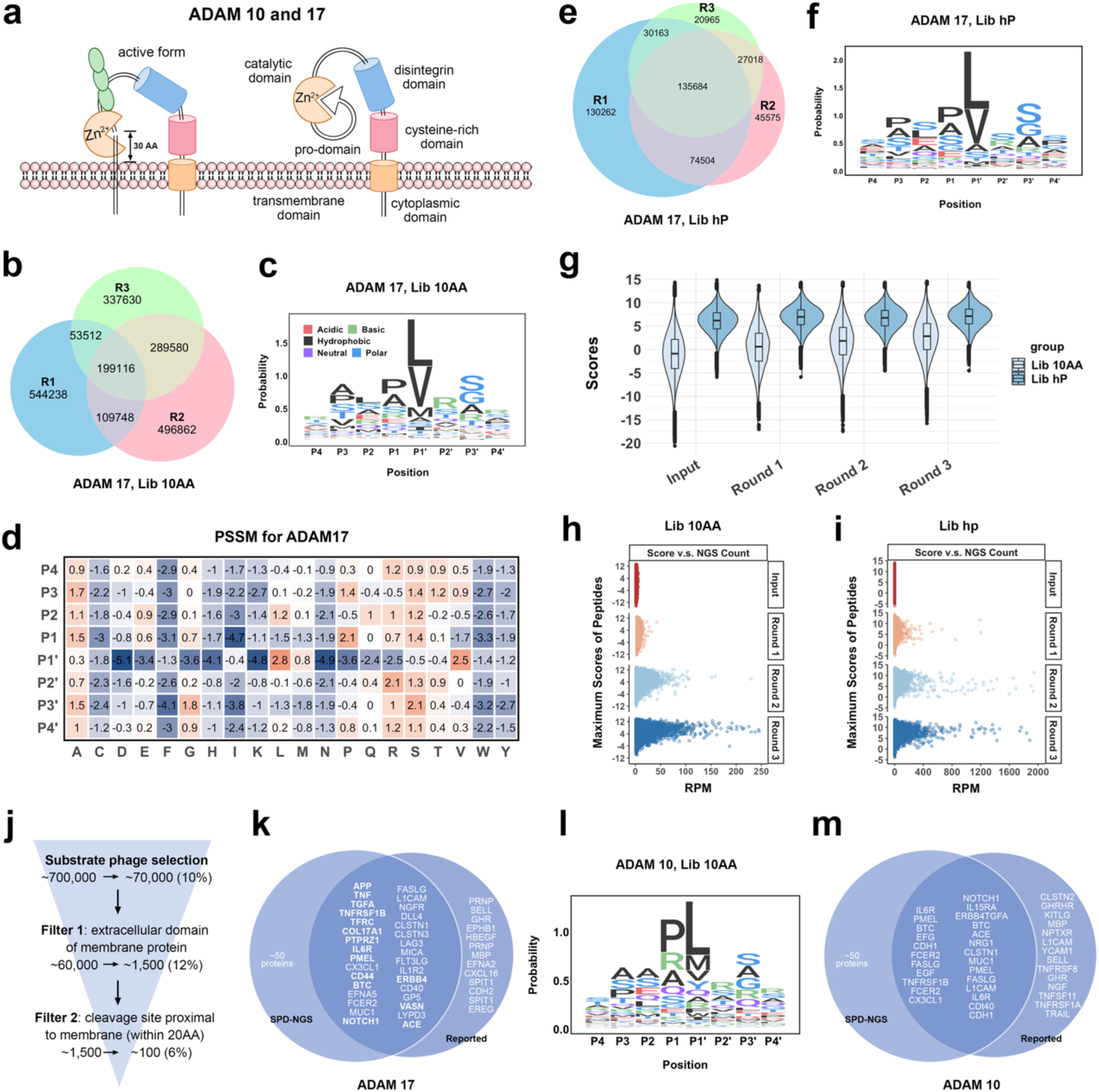
Mapping ADAM 10 and 17 proteases specificity. (**a**) Schematic illustration of ADAM 10 or 17 and the cleavage of their substrates proximal to the membrane. (**b**) The Venn diagram of individual peptides identified from Lib 10AA in each of the three rounds of the selection with ADAM17. (**c**) The sequence logo of ADAM 17 substrate consensus generated by aligning top 10,000 most frequently observed sequences from screening Lib 10AA. (**d**) The PSSM of ADAM 17 substrates generated according to the alignment. A score of every 8AA peptide window was calculated based on the PSSM. (**e**)The Venn diagram of unique peptides identified from Lib hP in each of the three rounds of the selection with ADAM 17. (**f**) The Sequence Logo of ADAM 17 substrates generated from the top 50,000 cleavage events identified from Lib hP. (**g**) Violin plot of the maximum scores of either Lib 10AA or Lib hP peptides in input library, Round 1, 2, and 3 outputs. The peptides with higher scores get enriched with more rounds of selection. (**h**) (**i**) The plot of maximum score vs NGS counts for Lib 10AA or Lib hP peptides, respectively, for each input library and Round 1, 2, 3 outputs. (**j**) The filters applied to identify protein substrates of ADAMs. (**k**) Venn diagram of ADAM 17 substrates identified in our screening and ones that have been reported in the literature (The substrates with a known cleavage site are in bold and details are in **Table 1**). (**l**) The Sequence Logo of ADAM 10 substrate consensus generated by 50,000 events identified from Lib 10AA. (**m**) Venn diagram of ADAM 10 substrates identified in our screening and the ones reported as a substrate (some of the substrates without a known cleavage site).

**Table 1.** ADAM17 cleavage sites in protein substrates reported before and identified by SPD-NGS. Reported cleavage sites are shown for proteins established as ADAM17 substrates through genetic, knockdown, or proteomics approaches. The same cleavage events were also identified in SPD-NGS with PSSM scores and their distance to membrane listed below. The distance to membrane was obtained from UniprotKB topological domain information.

| GENE | Cleavage Site | PSSM Score | Distance to Membrane |
| --- | --- | --- | --- |
| **TNF** | LAQAVRSS | 11.56 | 11 |
| **TGFA** | DLLAVVAA | 8.6 | 9 |
| **TNFR** | PAEGSTGD | 1.73 | 4 |
| **IL1R2** | EASSTFSW | 1.3 | -3 |
| **COL17A1** | EKDR LQGM | 2.26 | 35 |
| **TFRC** | ECERLAGT | 5.5 | 18 |
| **IL6R** | TSLPVQDS | 5 | 7 |
| **NOTCH1** | KIEAVQSE | 4.2 | 15 |
| **PTPRZ1** | LAEGLESE | 10.4 | 6 |
| **APP** | HHQKL VFF | <0 | 14 |
| **PMEL** | VSTQLIMP | 1.5 | 12 |
| **VASN** | TPPAVHSN | 2.21 | 17 |
| **CD44** | SHGSQEGG | 0.6 | 19 |
| **BTC** | DLFY LQQD | 1.8 | 7 |
| **ERBB4** | STLPQHAR | 5 | 6 |
| **ACE** | NSARSEGP | 3.2 | 24 |
| **TNFRSF1B** | FALPVGLI | 1.02 | 0 |

We applied our SPD-NGS libraries for deeper sequence coverage and to identify optimal sequences and potentially identify new protein targets. To identify the cleavage sites of ADAMs, we first expressed extracellular forms of ADAM 10 and 17 in human Expi293 cell (see Material and Methods). Briefly, after purifying them with Ni-NTA resin, we confirmed their activities by monitoring the cleavage of the known fluorogenic peptide substrate, Mca-KPLGL-Dpa-AR-NH2 (**Supplemental Figure 17**). After three rounds of phage selection against Lib 10AA using ADAM 17 (**Figure 5b, Supplemental Figure 18a)** we generated a sequence consensus from the NGS data set using MSA of the top ∼20,000 enriched sequences from Round 3 by fixing the large hydrophobic residue at P1’ (**Figure 5c**). Remarkably, the sequence logo was similar to that derived from ∼20 substrates from MEROPS sequence data (**Supplemental Figure 18b).** Based on the alignment we generated a PSSM over each the putative 8-mer substrate sequences from the Lib 10AA dataset (**Figure 5d)**. This showed a strong substrate preference for the hydrophobic residues, Val and Leu at P1’.

We next used the Lib hP to select for human derived sequences for ADAM17. After 3 rounds of selection (**Figure 5e**), we generated a sequence logo by aligning the top 20,000 sequences (**Figure 5f**). This largely matched the sequence logo derived from Lib 10AA (**Figure 5c**). We constructed violin plots of the maximum scores of the 10AA or 49AA sequences to monitor enrichment as a function of round of selection (**Figure 5g**). We see the change in the violin plots are more subtle than seen for the caspase selections (**Figure 2f, 2k**) which likely reflects lower specificity requirements for the ADAM17 substrates. The lower specificity of substrates for ADAM17 relative to caspases likely explains the larger number of substrates identified (**Figure 5b, 5e**). This is further supported by the lower raw NGS counts seen for top selectants as a function of round of selection for both Lib 10AA and Lib hP for ADAM17 (**Figure 5h, i**) compared to caspase-3 (**Figure 2g, l**) of 250+ RPM versus 1200+ RPM for ADAM17 and caspase-3, respectively.

The 10AA Lib results revealed a broad proteolytic specificity of ADAM17 for linear peptides. We next sought to identify human protein sequences. From Lib hP we found >100k substrates with >3NGS counts; this is clearly a gross over-estimate for what to expect for the roughly 50,000 protein isoforms in the human genome. To triage the candidate list of the ∼100k cleavable linear human sequences we applied two filters. First, we know that ADAM17 works on extracellular domains of membrane proteins; applying this criterion reduced the number of candidate sequences to roughly 1000 (**Figure 5j**). Proteomics data of the identified substrates show that cleavage occurs within a 30AA window beyond their transmembrane ecto-domains (**Figure 5a**, **Table 1**). The catalytic site in close proximity to the membrane-proximal cleavage sites of ADAM17 substrates.^34, 35^ This is reinforced by a recent X-ray structure of ADAM10, the closest relative of ADAM17, that has been solved.^33^ When we apply this structural filter for extracellular sequences within a 30AA juxta-membrane extracellular regions for Type I or II membrane proteins, we are left with about ∼100 putative substrates from the SPD-NGS data set (**Appendix Table S7**). Remarkably, this set captures virtually all of the annotated substrates for ADAM 17 that have been validated in the past two decades (**Figure 5k**, **Table 1**). The exact proteolytic sites for lots of the known protein substrates have not been validated yet for the reasons that i) ectodomain shedding occurs in the immediate extracellular juxtamembrane region, which is also where O-glycosylation is often found ii) multiple cleavages occur on the same stalk due to the low specificity and iii) the involvement of other proteases, like other ADAMs family proteases and peptidase. According to our PSSM, most 49ers enriched from SPD-NGS have multiple cleavage sites if we set threshold τ_i_ as 0, indicating the low specificity of ADAM17. In addition to Type I and II membrane proteins, we also identified cleavage events in proximal-membrane regions of extracellular domain of multi-pass transmembrane proteins (**Supplemental Figure 18c**) and GPI-anchored proteins. We are not aware of others reporting cleavage of multi-pass transmembrane protein by ADAM17 substrates because the fragments are not shed. We provide here a list of all predicted cleavages sites proximal to cell membrane (**Appendix Table S7**).

We applied the same workflow used for ADAM17 to ADAM10 its closest sequence relative (**Supplemental Figure 19, Appendix Table S8**). We conducted three rounds of selection with Lib 10AA, generated a sequence logo and PSSM from the MSA then applied these to Lib hP Round 3 selectants. This allowed us to generate a combined sequence logo shown in **Figure 5l** based on ∼100k selected sequences. We then triaged the ADAM10 substrate list from the Lib hP selectants based on sequences within 20AA of the juxtamembrane extracellular regions for Type I or II membrane proteins and identified ∼20 candidate sequences (**Figure 5m**). Remarkably, this analysis captured more than 2/3 of substrates reported in the literature or in MEROPS for ADAM10.

## Discussion

Substrate phage methods were developed more than two decades ago,^14^ but only recently have started to emerge as a useful tool for analysis of protease specificity.^16, 17, 33^ More recent advancements in the fields of synthetic DNA, NGS, bioinformatics now enable far deeper profiling of protease substrate specificity to identify consensus linear sequences and even candidate protein substrates at unprecedented and proteome-wide scales. SPD-NGS allows for deep profiling of protease specificity of linear peptides sequences at >1000 fold-depth over traditional natural peptide libraries. The sequence logos identified by SPD-NGS are in close agreement with literature curated proteomic data sets from the two classes we evaluated. In addition to the expanded sequence depth of coverage, there are several other important advantages. These libraries are genetically encoded so provide a simple, cheap, and renewable source accessible by simple molecular biology techniques. They are easily modified allowing synthesis of more focused libraries as illustrated in the sub-library built for the ADAMs proteases. These studies validate a broad framework for comprehensive protease profiling that complement and expand upon and existing synthetic peptide and proteomics technologies.

We found the randomized Lib 10AA to be useful to first calibrate cleavage conditions, and to develop consensus Logos over the typical 6-8 residue stretch that proteases engage. The depth of coverage allows one to analyze potential subsite cooperativity, a property that would be difficult to analyze with sparse substrate libraries. We analyzed this in the case of the caspases as interestingly found very weak subsite cooperative effects (**Supplemental Figure 20**). It was also possible to develop simple PSSMs based on multiple sequence alignments for both caspases and the less specific ADAMs proteases. This provided high confidence predictions for cleavage sites that matched literature expectations. The selections were highly reproducible and readily identified 20 to 100k unique cleavage sequences per substrate which is 100-1000-fold deeper coverage than typical synthetic peptide libraries. After three rounds of selection we find substrates that range over 1000-fold in NGS counts (ranging from 3 to 4000 NGS counts) demonstrating a broad dynamic range of substrate quality. Moreover, each of the five caspases and two ADAMs proteases that we tested had their own unique preferred sequence motif which highlights the sensitivity of the selections and the ability to generate highly specific protease fingerprints.

We show that the Lib hP to be useful for identifying candidate linear substrates in the human proteome. The PSSMs developed from the Lib 10AA were applied to the larger 49mer peptides in Lib hP to produce high confidence predictions for cleavage sites even when multiple cut sites were identified within the 49mer. The Logos generated from both the Lib 10AA and Lib hP were in close agreement and selections were highly reproducible. The Lib hP selections routinely captured 10 to 100-times more candidate substrates than those previously identified from a decade of proteomics studies; most of the substrates found from proteomics were contained within the Lib hP selected sets. However, not all were found in the SPD-NGS data which may reflect incomplete sampling in the library, or that some the reported proteomics data only specifies the protein target and not the site of proteolysis.

Although the Lib hP can broadly identify what can be cut not all will be accessible to the protease in the context of the folded protein. Thus, to help triage the candidate linear substrates, we found it useful to apply a global modeling and surface accessibility score, DSSP, to filter cleavage sites based upon surface accessibilities. The WARS protein served as one example. WARS had not been identified as a caspase substrate by proteomics but was found in the SPD-NGS screen for caspase-3. The DSSP analysis predicted two accessible sites, and both were confirmed by direct testing *in vitro* and in the same rank order as predicted by DSSP analysis. Secondly, it is useful to triage substrates based upon the cellular location of the protease. In the case of the intracellular caspases one can exclude targets that are extracellular or within vesicles. This was especially useful in the case of the ADAMs proteases for triaging 60k candidate 49 mer substrates that were cleaved by the ectodomain of the two ADAMs proteases. These candidate substrates triaged to 1k when considering only domains that were extracellular and to 30 when considering the structural restrictions that ADAMs cleave substrates within 20 AA of the nearest transmembrane ecto domain. Indeed, this triage approach captured virtually all of the known substrates and for both of the ADAMs.

We applied a two-library approach when deriving PSSMs. Data from the Lib 10AA was critical for deriving a PSSM for the caspases and ADAMs that could then be applied to the larger 49mers in the Lib hP. The reason for these two steps is that the PSSM requires a multiple sequence alignment which is accessible within 10mers but beyond reach in larger 49mers.

Although SPD-NGS from Lib hP identifies far more candidate human substrates than are likely present in folded proteins, the data can be informatically triaged by compartment and structural considerations as described above. Moreover, we believe the data could be a very useful companion for targeted mass spectrometry experiments by providing look-up lists for parallel reaction monitoring.^11, 19^ The SPD-NGS approach based on linear peptides does not take into account the possibility of distal exo-sites that have been seen in rare cases for caspase-7.^36, 37^ One way to expand the system to native proteins would be to display whole proteome libraries of folded proteins that may be in reach given the low cost of synthetic DNA and orfeome cDNA libraries. Another limitation is that the system is currently restricted to screening purified endo-proteases, not exo-proteases because the N- and C-terminus is blocked on the phage. Despite these current limitations, we believe SPD-NGS is now a useful first pass companion to more traditional peptide library and proteomics methods given the simplicity of the selections and analysis, renewability, reproducibility, depth of coverage, speed, and low cost.

## Materials and Methods

### Protein expression and purification

ΔCARD-caspase-2, caspase-3, 6, 7, ΔDED-caspase-8 and WARS were cloned into His6-affinity tag containing vector pET23b. The active enzymes were expressed in *E. coli* BL21 (DE3) pLysS cells (Promega, # L1195) and WARS was expressed in *E. coli* strain BL21(DE3). Cells were grown in 2xYT media supplemented with 200 μg/mL ampicillin and 50 µg/mL chloramphenicol (for pLysS only) at 37 °C to an OD600nm at approximately ∼0.6–0.8. Expression was then induced for the caspases and WARS with IPTG (details of concentration, temperature, and duration are available in table below). Cells were harvested by centrifugation and lysed by sonication. The cell lysates were clarified by centrifugation, and the soluble protein fractions were purified by Ni-NTA resin (Qiagen, # 30230). The purified fraction was concentrated, and stored at −80 °C.

| Protein | IPTG (mM) | Temperature (°C) | Time (hour) |
| --- | --- | --- | --- |
| ΔCARD-caspase-2 | 0.2 | 30 | 4-6 |
| caspase-3 | 0.2 | 30 | 4-6 |
| caspase-6 | 0.05 | 30 | 16 |
| caspase-7 | 0.2 | 30 | 16 |
| ΔDED-caspase-8 | 0.4 | 30 | 3-5 |
| WARS | 0.2 | 18 | 16 |

The vector used to express ADAM10, 17 ECD-10His (extracellular domain) was generated by Gibson cloning and adapted from the commercially available pFUSE-hIgG1-Fc (InvivoGen) vector. Each ECD was subcloned between an N-terminal IL2 signal sequence and a C-terminal 10His-tag replacing the original Fc domain. Suspension Expi293 cells were grown to 2.5 million cell density and transiently transfected with ADAMs protease expression vectors using ExpiFectamine™ 293 Transfection Kit (Thermo Fisher Scientific, # A14525). Medium was harvested after 6-7 days and protein was purified by Ni-NTA chromatography and assessed for quality and activity.

### Phagemid construction

The pKM128 vector (phagemid) from Kurt Mou was modified to parental vectors, PhMd0007 and PhMd0003, for Lib10AA and Lib hP, respectively. An Avi tag was first inserted into the phagemid after a PhoA promoter, followed by a TAA stop codon, an EcoRI restriction site, a TEV cleavage site and the truncated gIII protein for PhMd0007 (Lib 10AA). For Lib hP, PhMd0003 has an Avi tag, followed by an EcoRI restriction site, a BamHI restriction site, a TAA stop codon, a HindIII restriction site, followed by the truncated gIII. All oligonucleotides for cloning were from Integrated DNA Technologies.

### Construction and characterization of Lib 10AA and Lib hP

The ten-amino acid randomized linker in Lib 10AA was constructed using synthetic DNA containing 32 NNK degenerate codons (where N=A/G/C/T and K=T/G) encoding all 20 amino acids at each of the 10 positions (**Supplemental Figure 1**). The synthetic DNA was incorporated into the phagemid template by Kunkel mutagenesis (**Supplemental Figure 1a**).^24, 38^ The theoretical sequence diversity of Lib 10AA is ∼10^13^ but the true diversity was titered after transfection at ∼10^9^. NGS sequencing of the unselected library confirmed the broad representation of sequences in the library and revealed an amino acid or nucleotide distribution very close to the theoretical values based on the input synthetic DNA **(Supplemental Figure 1b, 1d**). The NGS also showed the amino acid and nucleotide diversity was uniform across all 10 positions as programmed by the synthetic DNA **(Supplemental Figure 1c, 1e**).

The Lib hP was generated by subcloning from a human tiled T7 phage library in 49AA blocks with 25A overlaps as previously described^39^ (**Supplementary Figure 2**). Briefly, the synthetic DNA for the T7 library was generated from all human sequences (variants and isoforms) in the NCBI protein database (Nov 2015) and redundancy collapsed based on 95% sequence identity. These were split into 49-residue blocks and tiled in 25-residue overlaps to reduce the risk of losing library diversity and to provide duplicate coverage. This yielded a total of 731,000 sequences that were codon optimized for expression in *E. coli*. We PCR amplified the oligos pools for T7 library provided by Prof. DeRisi Lab with proper flanking region and cloned into our M13-phagemid to generate Lib hP. The quality of Lib hP was evaluated by NGS (150nt single reads on Ilumina HiSeq4000 or NextSeq). The NGS data of the starting oligo pools revealed that 93% of the sequences were human derived and error-free whereas 89% of Lib hP was human derived and error free (**Supplemental Figure 2b**). This indicated the subcloning step retained virtually the same diversity and quality as the starting library. Lib hP had and 6% had deletion and 1% have insertion while 88% of total sequences could be detected by NGS in the constructed library, among which 89% are error-free, 9% have deletion and 1% have insertions. Considering the 25-residue overlap and NGS errors, we would expect Lib hP to have higher sequence coverage. As expected, the amino acid composition was very uniform across the 49 positions except for the enrichment of methionine at the first position that was programed as a start codon for protein translation from the T7 library (**Supplemental Figure 2c, 2d**).

The tiled peptide library (Lib hP) covers the entire human proteome, thus should give us more comprehensive profile than with proteomics because no cell line expresses all the proteins at one time. A single bulk cell proteomics experiment or RNAseq from any given cell line would only identify 8,000-10,000 proteins or 10,000-15,000 transcripts, respectively (**Supplementary Figure 2e**). Using RNAseq, the minimum numbers of cell lines that are required to achieve different gene coverage is shown in **Supplementary Fig. 2f.** One can estimate for 95% gene coverage would require more than 40 cell lines.

### Construction of randomized 10AA synthetic substrate phage library (Lib 10AA)

#### Isolating ssDNA of phagemid

The parental phagemid was first constructed in which an avi-tag was linked directly to a sequence including an EcoRI restriction site, a TAA stop codon, followed by a truncated M13 gene III protein. After transformation, a single colony of *E. coli* CJ236 (dut^-^/ung^-^) was picked and grown in 1 mL 2XYT supplemented with appropriate antibiotics. After the culture reached O.D. 0.6, M13K07 helper phage (10^10^ PFU ml^-1^) was added and the culture was incubated at 37 °C, 250 RPM. After 1 hour, the culture was transferred into 500 mL 2XYT supplemented with appropriate antibiotics, uridine and grown overnight. The next day, phage particles were precipitated with 1/5 volume of PEG/NaCl solution and collected by centrifugation. The phage pellet was resuspended in PBS buffer and ssDNA was purified using a MiniPrep column. Briefly, phage solution was applied to MiniPrep column to bound the phage to the column matrix and 0.7 mL of buffer MLB to the column twice to lyse the phage particles. The ssDNA on the column was washed by 0.7 mL PE buffer twice and eluted by ddH_2_O.

#### Kunkel mutagenesis to generate heteroduplex CCC-dsDNA

The first step in the synthesis of heteroduplex CCC-dsDNA is the phosphorylation of the mutagenic oligonucleotide. The mutagenic oligonucleotide was 5’-phosphorylated to enable ligation by T4 DNA ligase at 37 °C for 1 hour.

| Component | Volume (μL) | Final Concentration |
| --- | --- | --- |
| Oligonucleotide (100 uM) | 3 | 9 ug |
| 10 X PNK Reaction Buffer | 5 | 1X |
| ATP, 10 mM | 5 | 1 mM |
| T4 PNK (10 U/uL) | 5 | 50 U |
| ddH<sub>2</sub>O | 32 | 50 uL |

After phosphorylation, the oligos were annealed to the ssDNA as following:

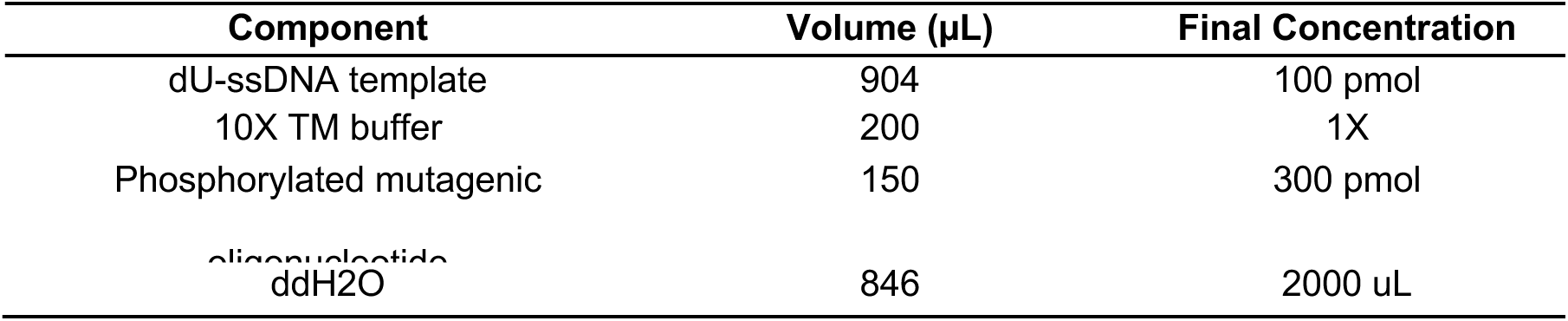

The above annealing reaction mixture was prepared in a 1.5-ml microcentrifuge tube and aliquoted into 50 PCR tubes. The annealing reaction was conducted at 90 °C for 3 min, 65 °C for 5 min, 63 °C for 5 min, 61 °C for 5 min, 59 °C for 5 min, 57 °C for 5 min, 20 °C for 5 min. To complete the enzymatic synthesis of CCC-dsDNA, 6.5 µL of the following solution was added to each of the 50 reactions above. The reaction was completed overnight at 20 °C.

| Component | Amount (μL) | Final |
| --- | --- | --- |
| ATP, 10 mM | 100 | 346 mM |
| dNTP mix (25 mM of each) | 100 | 865 mM of each nucleotide |
| DTT, 1M | 15 | 5.19 mM |
| T4 DNA ligase | 30 | 2,000 ligation units or 30 Weiss units |
| T7 DNA polymerase (10 | 30 | 30U |

The ccc-dsDNA was purified by PCR purification column. To improve electroporation efficiency, drop dialysis was conducted to remove excess salt in the DNA. The DNA can be frozen at -20 °C for later use.

#### E. coli electroporation and phage propagation

A 1-mm gap electroporation cuvette used in electroporation was pre-chilled on ice. A 50 µL aliquot of electrocompetent *E. coli* SS320 was thawed and mixed with 5 µL DNA before being transferred into the cuvette. Electroporation was performed following the manufacturer’s instruction with a Bio-rad Gene Pulser with the following settings: 1.0 mm cuvette, 10 µF, 600 Ohms, 1800 Volts. The electroporated cells were immediately rescued by adding 1 mL prewarmed SOC and transferred to 10 mL SOC media in a baffled flask. The cuvette was rinsed twice with 1 mL SOC media. A serial dilution was made to determine the library diversity (typically 1-3 x 10^9^). After 30-min recovery, all the SOC culture was transferred into 1 L 2XYT/carb/kan/M13K07 helper phage media and shaken at 250 rpm at 37 °C for 20 hours.

The next day, the culture was centrifuged and the supernatant was collected. Phage were precipitated by adding 1/5 volume 5X PEG/NaCl precipitation buffer (20% PEG-8000, 2.5 M NaCl) and centrifuged at 9,000 RPM for 20 min. Pellets were resuspended in ¼ starting volume storage buffer (1 X PBS, 0.05% tween-20, 0.2% BSA) supplemented with protease inhibitor. The library was stored at -80 °C freezer supplemented 10% glycerol.

### Construction of human proteome substrate phage library (Lib hP)

#### Vector digestion

The parental phagemid was first constructed in which an avi-tag was linked directly to a sequence including an EcoRI restriction site, a TAA stop codon, a BamHI restriction site, and a HindIII restriction site, followed by a truncated M13 gene III protein (**Figure 1a**). The EcoRI and HindIII restriction sites was for cloning and the middle BamHI site was designed for removing parental phagemid in the cloning process. The parental phagemid was then subjected to restriction digestion under the following reaction conditions:

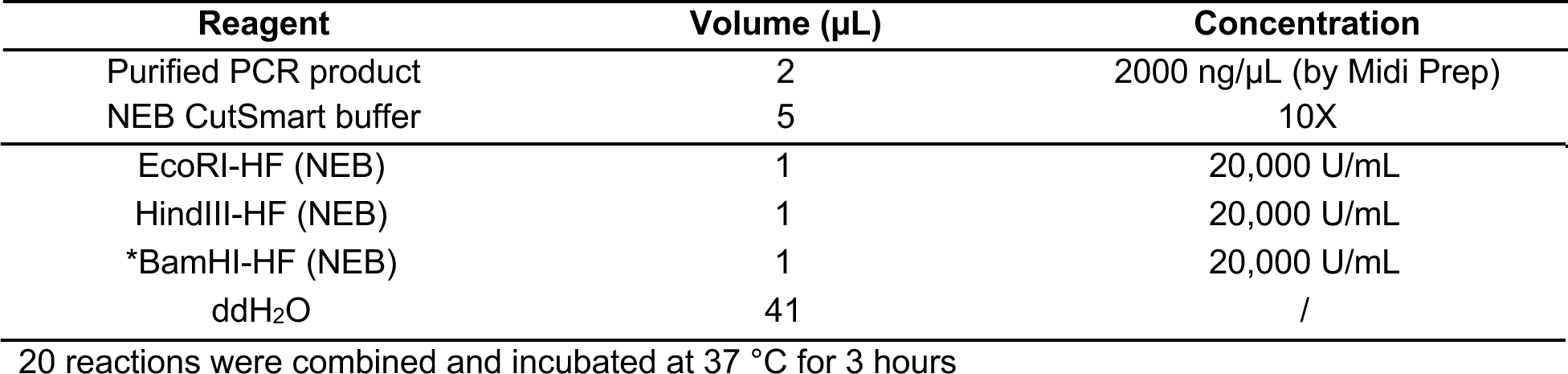

| Reagent | Volume (µL) | Concentration |
| --- | --- | --- |
| Purified PCR product | 2 | 2000 ng/µL (by Midi Prep) |
| NEB CutSmart buffer | 5 | 10X |
| EcoRI-HF (NEB) | 1 | 20,000 U/mL |
| HindIII-HF (NEB) | 1 | 20,000 U/mL |
| *BamHI-HF (NEB) | 1 | 20,000 U/mL |
| ddH<sub>2</sub>O | 41 | / |

20 reactions were combined and incubated at 37 °C for 3 hours

After digestion, the digested vector was size selected by agarose gel electrophoresis and concentrated by GeneVac before ligation/cloning.

#### Insert preparation

The single-stranded DNA oligonucleotide library was generously provided by DeRisi Lab at UCSF. Oligos were amplified by PCR with primers adding relevant EcoRI and HindIII restriction sites. 1 µL of 5 nM oligo is used for 96 25-µL PCR reactions:

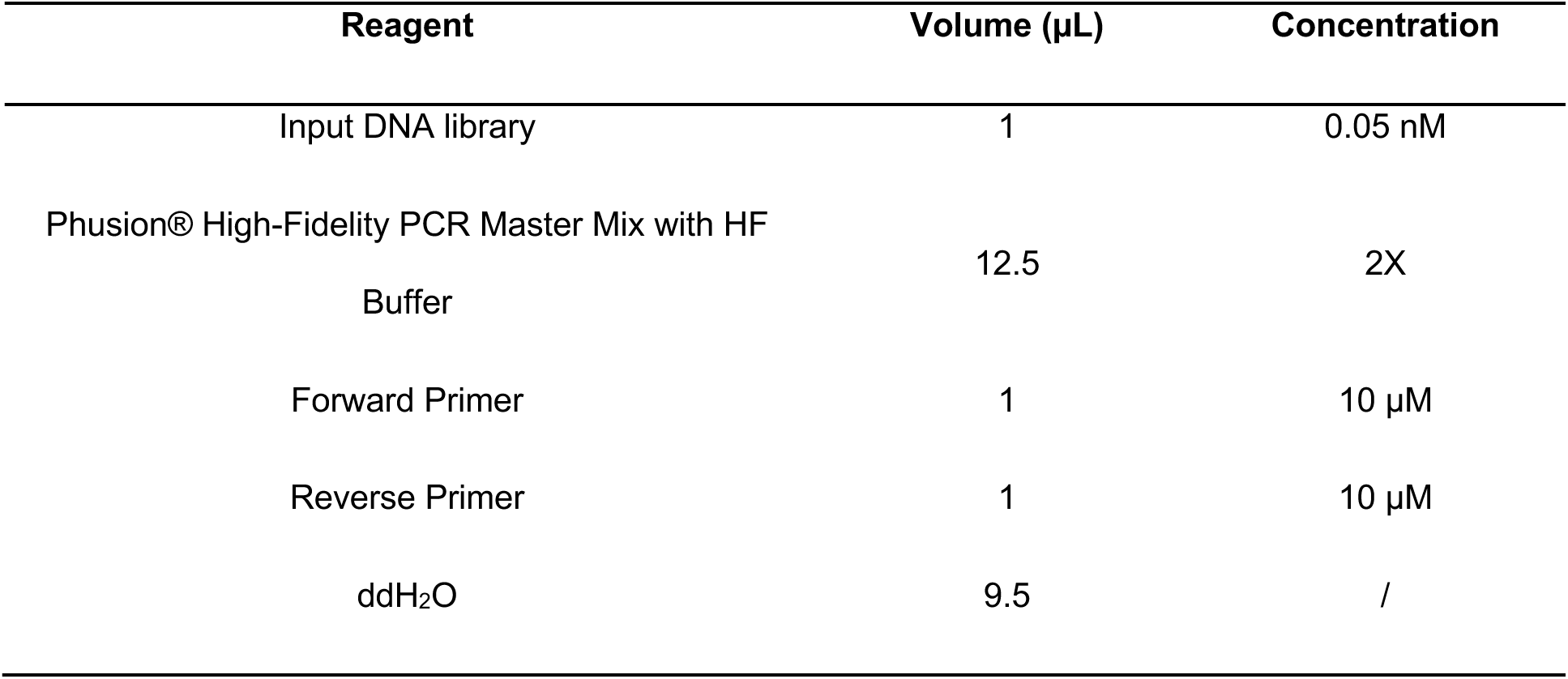

#### Primer design

Forward Primer: ATGGCATGAA<u>GAATTC</u>TGGAGCCATCCGCAGTTCG (underscored: EcoRI restriction site)

Reverse Primer: AATCAAAATC<u>AAGCTT</u>CTTATCATCGTCGTCCTTGTAGTC (underscored: HindIII restriction site)

Thermocycler setup:

| Temperature (°C) | Times (mm:ss) | cycles |
| --- | --- | --- |
| 98 | 03:00 | 1 |
| 98 | 00:30 | 25 |
| 72 | 00:30 |  |
| 72 | 10:00 | 1 |
| 4 | ∞ | / |

All the PCR products were combined and purified by AMPure PCR purification magnetic beads. DNA was then subject to restriction digestion under the following reaction conditions:

| Reagent | Volume (μL) | Concentration |
| --- | --- | --- |
| Purified PCR product | 10 | 300 ng/μL |
| NEB CutSmart buffer | 5 | 10X |
| EcoRI-HF (NEB) | 1 | 20,000 U/mL |
| HindIII-HF (NEB) | 1 | 20,000 U/mL |
| ddH<sub>2</sub>O | 33 | / |

Reaction was incubated at 37 °C for 2 hours

The digestion mixture was purified by Qiagen PCR purification kit. The sizes of the fragments were confirmed by agarose gel electrophoresis before cloning.

#### Cloning/Ligation

The ligation reaction was set up in the PCR tubes according to the table below:

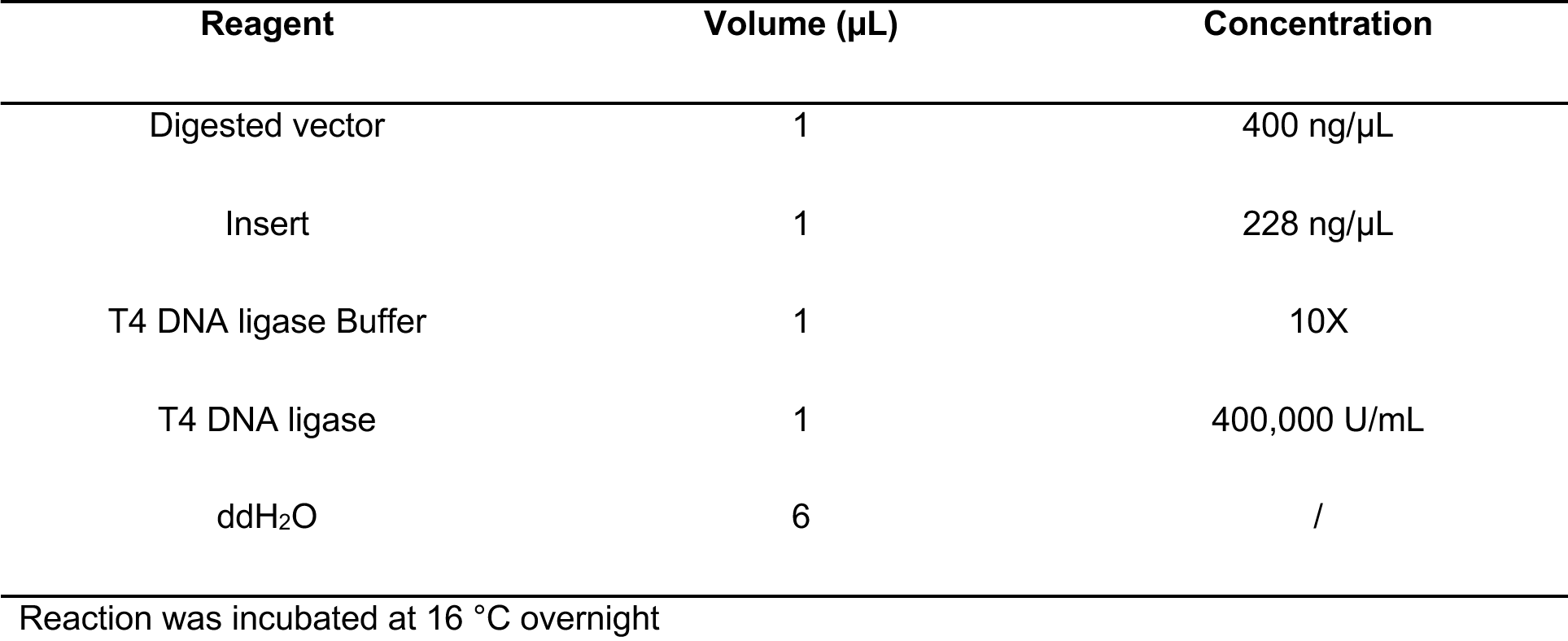

The ligation mixture was purified by Qiagen PCR purification kit and then concentrated by GeneVac. To improve electroporation efficiency, drop dialysis was conducted to remove excess salt in the DNA. The DNA can be frozen at -20 °C for later use.

### In vitro biotinylation of the phage library with BirA ligase

10 times more BirA ligase was used for the biotinylation reaction than a typical condition because BirA activity was partially inhibited by 10% glycerol. Generally, 120 µL or BioMix A (10X concentration: 0.5M bicine buffer, pH 8.3), 120 µL of BioMix B (10X concentration: 100mM ATP, 100mM MgOAc, 500μM d-biotin)), and 10 µL (10 mg/mL) BirA ligases were added into 1 mL substrate phage library with an Avi-tag on the N-terminal of the displayed peptides. The reaction mixture was incubated at 37 °C for 1 hour. The phage were then precipitated in 30 mL PBS buffer supplemented with 0.05% Tween20 and 0.2% BSA (PBSTB) and 1/5 volume 5X PEG/NaCl. The phages were pelleted with centrifuge at 9000 RPM, 20 min and the supernatant was completely removed. To move extra free d-biotin, the phage were resuspended in PBSTB 1mL, precipitated with 1/5 volume 5X PEG/NaCl, pelleted by centrifuge twice.

### XL-1Blue BirA strain construction

To facilitate the screening, we have engineered the XL-1 Blue cells to express the biotin ligase BirA with pBirAcm, an engineered pACYC184 plasmid with an IPTG inducible birA gene. Briefly, competent XL-1Blue cells were transformed with pACYC184 plasmid harboring chloramphenicol resistance. On the day prior to round 1 selection, a single colony of XL-1 Blue BirA was inoculated into 100 µL 2XYT. The 100 µL culture was then split into 3 2-mL 2XYT cultures containing tetracycline (5 µL/mL)/chloramphenicol, carbenicillin (50 µL/mL), or kanomycin (25 µL/mL) in a 12mL falcon *E coli* culture tubes. These tubes were incubated in a 37°C, 250 RMP shaking incubator overnight to allow tetracycline/chloramphenicol resistant clones to grow out.

### General phage selection protocol

To perform a screening, 1 mL of biotinylated substrate phage library was incubated with 100 µL magnetic StreptAvidin (SA) beads for 30 min and after that, the SA beads were then stringently washed with PBS buffer supplemented with 0.05% Tween20 and 0.2% BSA to get rid of any free phage. After the SA beads were treated with protease of interest, released phage in supernatant were collected and propagated as protease-sensitive pool, and those bound on the beads can also be propagated as protease-resistant pool after released by non-specific proteases like trypsin. After three rounds of selection, phage polymerase chain reaction (PCR) was conducted, followed by NGS to identify sensitive and resistant substrate sequences in each pool. We conducted duplicates to assess reproducibility. After sequencing, we tally the read count for each peptide in both sensitive and resistant pools and process the data by a custom R script.

### NGS sample preparation

Propagating the phage mixture to saturation in culture followed by PCR was more accurate than direct PCR without propagation which required more PCR cycles.^40^ We thus used overnight cultures for NGS sample preparation after boiling the phage to extract ssDNA as templates for PCR reactions. For all samples from both Lib 10AA and Lib hP, a barcoding PCR process was performed using primers listed in **Supplemental Figure 21.** The thermal PCR profile for Lib 10AA was: 98 °C (20 s), 63 °C (15 s), 72 °C (30 s), 15 cycle. The thermal profile for Lib hP was: 98 °C (30 s), 72 °C (30 s), 15 cycle to prevent heterodimerization due to the 24-residue overlap. The number of cycles was determined empirically to prevent product laddering, assessed by agarose gel electrophoresis. PCR products were gel purified on 1.2 % agarose. Illumina library quality was assessed by Agilent DNA 1000 Bioanalyzer kit (5067-1504, Agilent), according to manufacturer’s instructions. Libraries were sequenced on a NextSeq or HighSeq 4000 (Illumina) using single-read 50 or 150 base pair reads. A custom sequencing primer was used (order as shown):

ATCGGATCCGCCGCTCTGAAAGTACAGATTCTC (reverse, Tm =67 ◦C, GC% = 52) or CTGCAGTATGTGGCATGAATCTGGTGGTGGT (reverse, Tm =67 ◦C, GC% = 52) for Lib 10AA and GTATGACGTCGCCATCCGCAGTTCGAGAAA (forward, Tm =67 ◦C, GC% = 52) for Lib hp.

### Sequencing analysis pipeline

Sequence filtering and peptide analysis were performed using an in-house informatics pipeline written mainly in R. Sample scripts are available for download (https://github.com/crystaljie/NGS_data_process_sample_script_for_substrate_phage_paper_JZHOU.git). The initial data processing for Lib 10AA and Lib hP are different. Raw NGS data from Lib 10AA, “*.fastq.gz” sequencing files were converted into a table with DNA sequences, amino acid sequences, counts/frequency, RPM as four columns, which were saved as .csv files for further analysis. For Lib hP, reads were aligned to a reference dataset using bowtie2 to generate Sam files. Sam files were then converted into Bam files using Samtools, which were then parsed using a suite of in-house analysis tools (R) to make a table with gene, ref.name, individual phage counts/frequency, peptide sequence, etc. as columns. These tables are saved as .csv files for further analysis. Some quality filters were applied: i) all the sequences with a stop codon are removed (Lib 10AA); ii) only the sequences that were in-frame are kept (Lib 10AA); iii) the bowtie2 alignment mapping quality should be greater than 10. Although we applied this stringent cut-off, it does not necessarily mean sequences that showed up only once in NGS may not be a possible substrate.

### Generation of PSSM

The first step towards building Scoring Matrix for predicting proteolytic sites involves an unbiased sequence alignment and we illustrate it with the caspase-3 data (**Supplementary Figure 8**). For practical reasons of computing time we aligned the top 20,000 most enriched sequences in Round 3 from Lib 10AA (**Supplementary Figure 8a**) to generate a position frequency matrix (PFM), the most basic representation of a motif and simply for each position the total counts of each amino acid (*C_N_*) (**Supplementary Fig. 8b**). 20,000 sequences guarantee the variability of the collection of proteolytic sites, which enhances the accuracy for predicting additional sites. Based on the alignment, the position frequency matrix (PFM) was generated, converted to a position probability matrix (PPM, **Supplementary Figure 8c**) and then position weight matrix (PWM) using formula shown below, where *B_N_*, a background probability for amino acid is 0.05 (**Supplementary Figure 8d**).

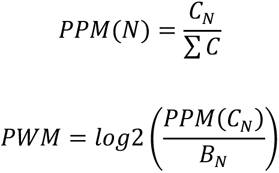

PWMs are also known as position-specific scoring matrices (PSSMs). Since starting from such a large pool of sequences, all the amino acids are scored in the matrix and there no need to use a pseudocount. Using this equation, the log of fractions where the probability of a certain amino acid in a sequence is higher than that of the background probability of that amino acid result in positive scores, and vice versa for negative scores.

### Data Archival

The sequencing data that support the findings of this study are available in the Gene Expression Omnibus (GEO) with the identifier ?. All other data supporting the findings of this study are available within the Appendix Table S1-S8.

## Acknowledgments

We thank the members of the Wells laboratory and Antibiome Center for inspiring and helpful discussions. We thank M. Hornsby, X. Zhou, and M. Zhong for sharing tips and tricks to construct phage libraries with Kunkel mutagenesis, B. O’Donovan, M. Raghavan, and C. Mandel-Brehm for suggestions on constructing the human proteome library, X. Zhou and J. Glasgow for inspiring discussion on how to optimize the experiments, X. Zhou for sharing pKM0128 phagemid vector, K. Leung for help on R programming, Y. Yang for coding in bash. J.A.W. thanks The Chan Zuckerberg Initiative and Biohub Investigator Program as well as NCI grant P41CA196276 for financial support of this work. J.Z. was supported by a NIH postdoc fellow F32CA236151-02. J.Y.Z. is supported by NSF CCF 1763191, NIH R21 MD012867-01, NIH P30AG059307, and grants from the Silicon Valley Foundation and the Chan-Zuckerberg Initiative.

## Author Contributions

J.Z. and J.A.W. designed the research. J.Z. performed all the experiments. J.Z. K.L. and J.A.W. developed the PSSM for data analysis and interpretation. S.L. and J.Y.Z. analyzed data using machine learning method. B.O. and J.A.D designed and ordered the human proteome oligo for library construction. J.Z. and J.A.W. wrote the manuscript and all provided editorial comments.

## Supplementary Information

**Supplementary Figure 1.**
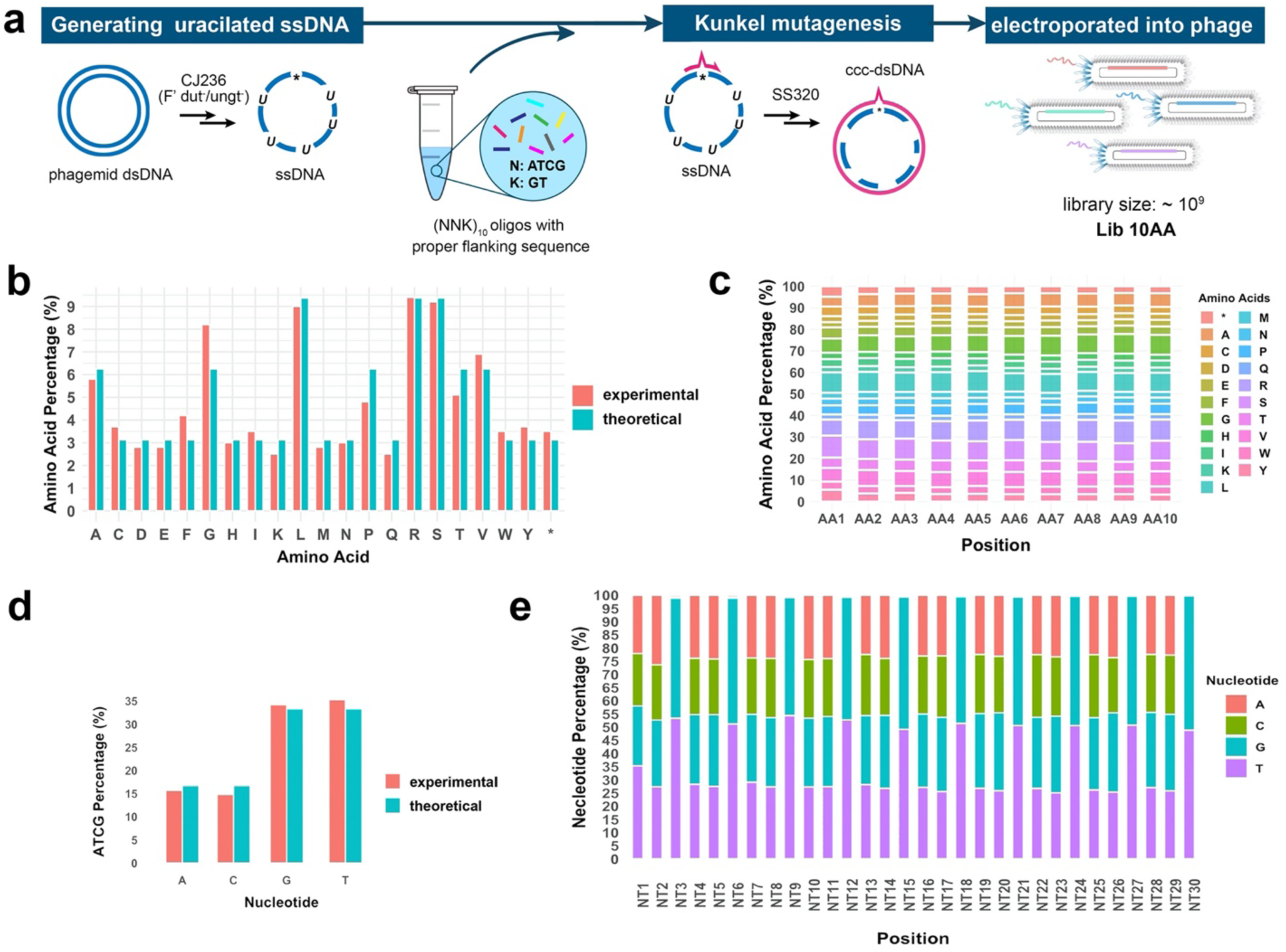
The construction and characterization of Lib 10AA for identifying protease substrate specificity. (a) Kunkel mutagenesis was used to construct Lib 10AA, that displays randomized 10AA peptides encoded by NNK codons. N encodes A, T, C, G while K encodes T and G, thus encoding 32 different tri-nucleotides encompassing all 20 amino acids. (b) The amino acid abundancy of Lib 10AA assayed by NGS closely matches the theoretical values (3.33%, 6.66% and 9.99%, depending on amino acid code degeneracy). (c) The amino acid composition established by NGS is calculated for each of the 10 positions in Lib 10AA and found to closely match the synthetic oligonucleotide input. (d, e) The nucleotide abundancy of Lib 10AA assayed by NGS closely matches the theoretical values and NNK pattern.

**Supplementary Figure 2.**
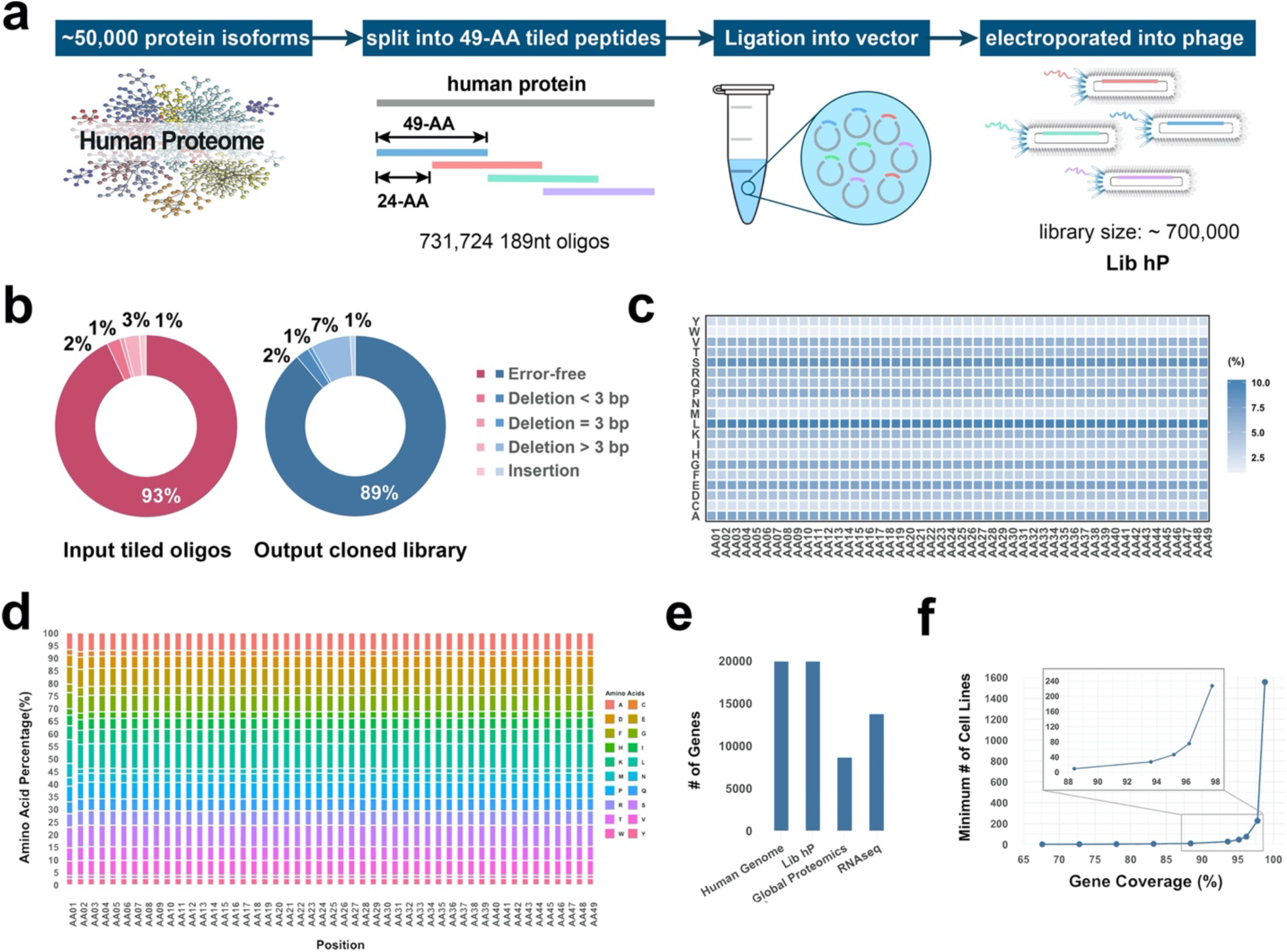
The construction and characterization of Lib hP for identifying protease substrate specificity. (a) Construction of the Lib hP, that displays tiled peptides covering the human proteome in 49AA blocks. (b) Assessment of the library quality before cloning and after library generation of Lib-hP by NGS shows high recovery of sequences. (c, d) The amino acid composition calculated for each of the 49 positions in Lib hP is representative of the composition found in the human proteome. (e) Protein expression gene coverage of Lib-hP, a single global proteomics experiment, and a single RNA-seq experiment. e) Minimum numbers of cell lines that are required to achieve different gene coverage in RNA-seq. Data analysis was based on RNA-seq of 1556 cell lines (NIH RNA-seq of 934 human cancer cell lines from the Cancer Cell Line Encyclopedia*^1^* and Genentech RNA-seq of 675 commonly used human cancer cell lines*^2^*).

**Supplementary Figure 3.**
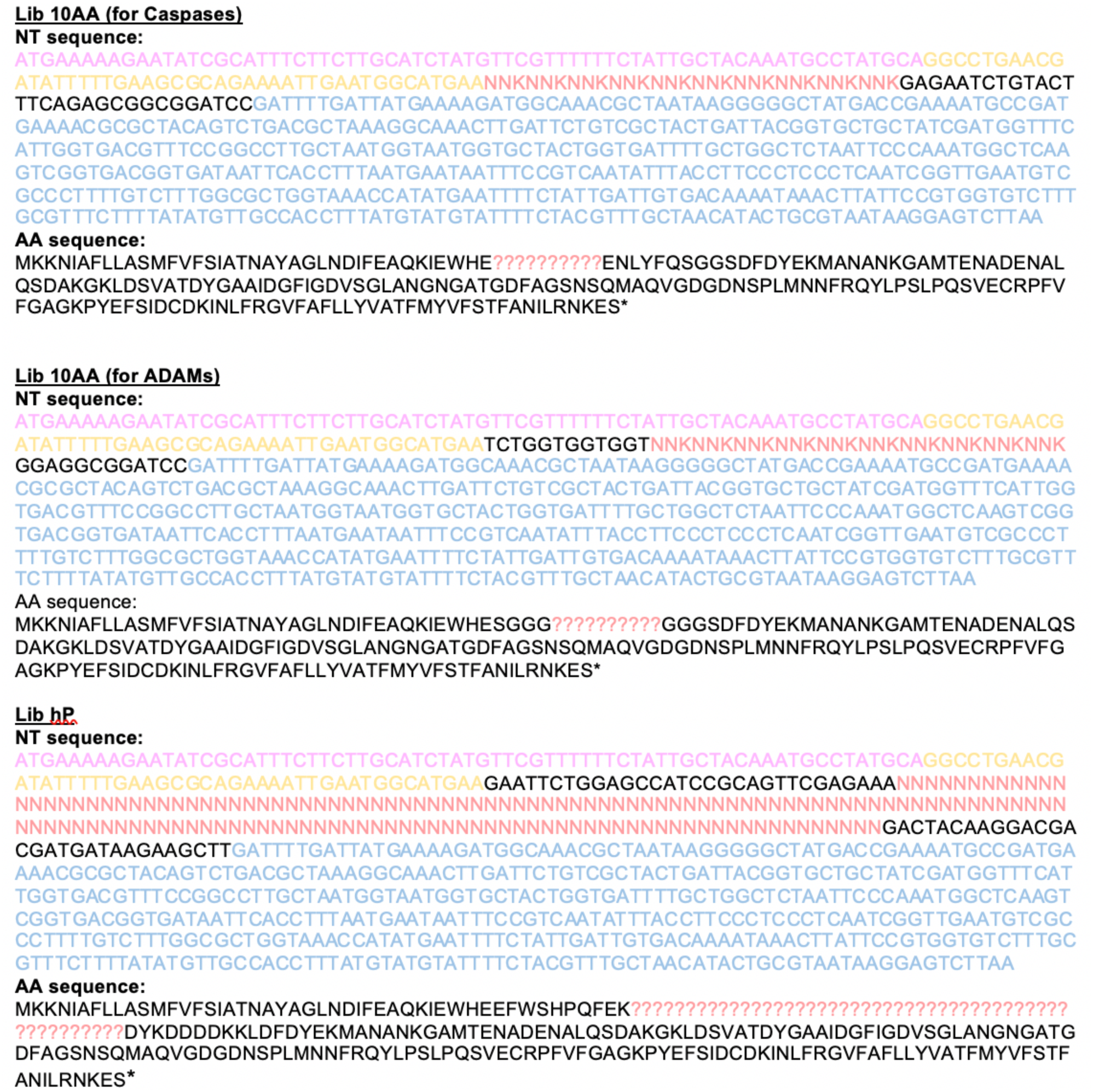
Phagemid DNA and protein sequences used for phage library construction.

**Supplementary Figure 4.**
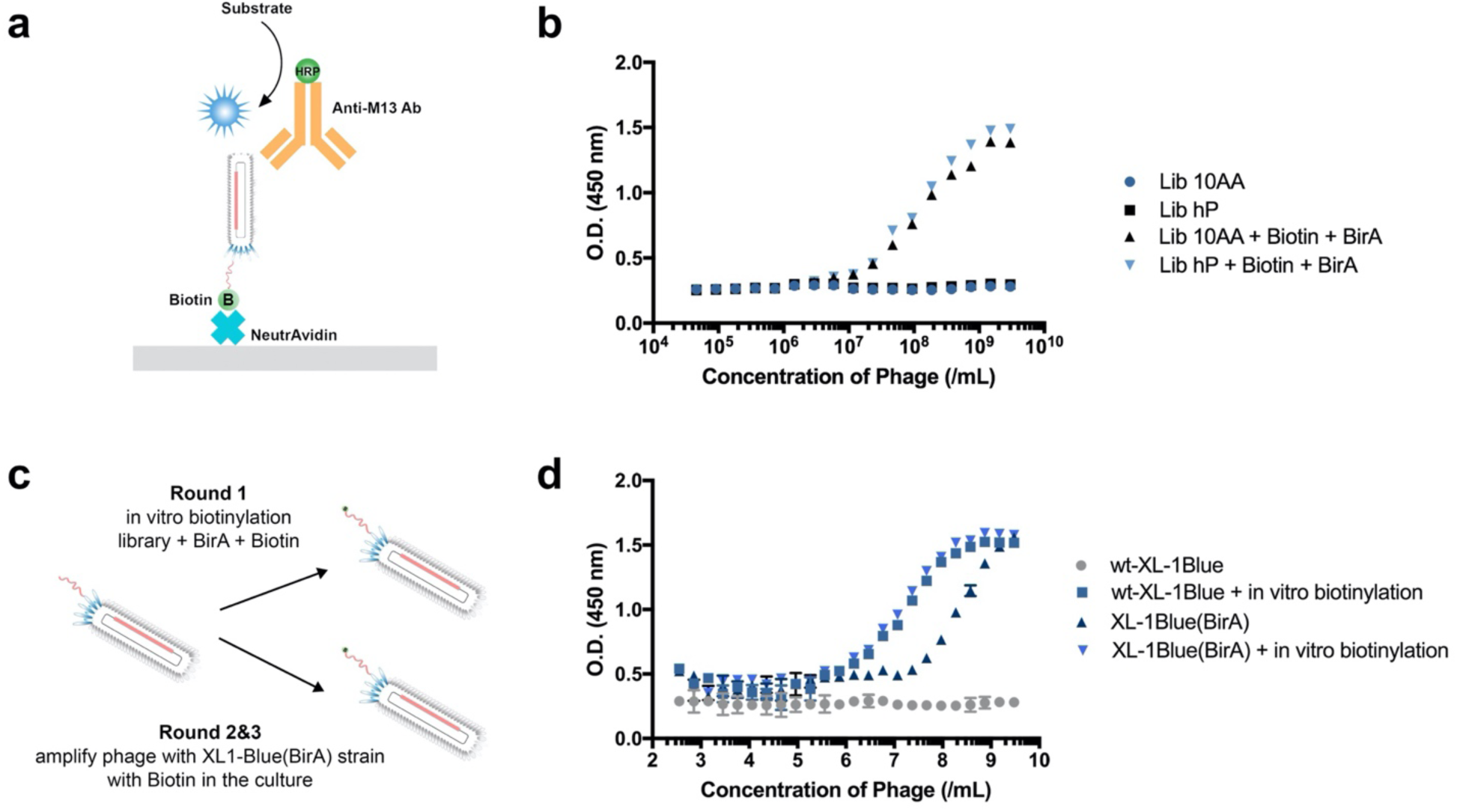
Biotinylation of the library before screening. (a) The experiment setup of phage ELISA for quantifying biotinylation level. (b) Phage ELISA confirmed that peptides of Lib 10AA and Lib hP were successfully displayed. The entire library (1 mL) was biotinylated by 5 µL, 10 mg/mL BirA in buffer with 100 µM D-biotin. (c) The library was first biotionylated with BirA in 100 µM D-biotin before Round 1 selection, while for Round 2&3 phage selection, biotinylation was done during phage amplification using engineered XL1-Blue (BirA) strain. (d) In cell biotinylation was doable although not 100% complete.

**Supplementary Figure 5.**
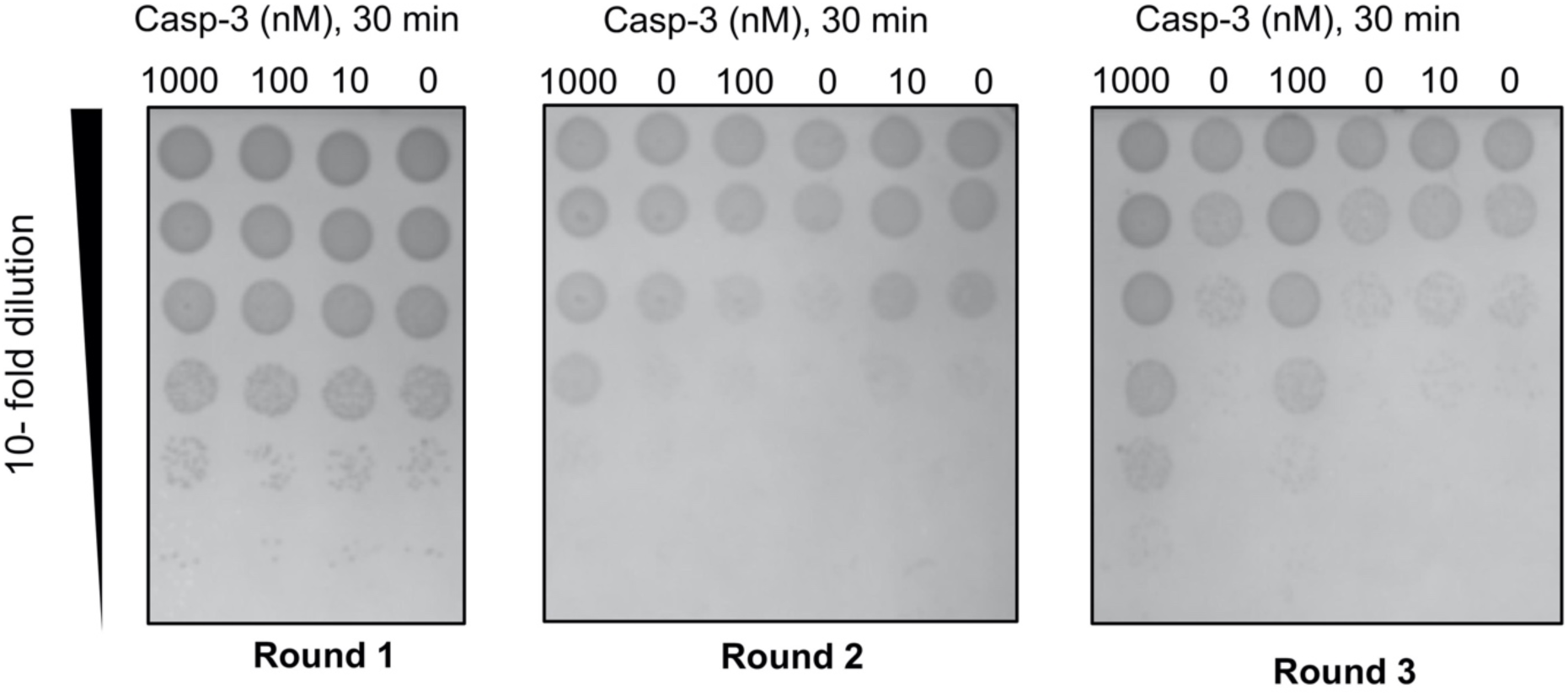
Phage titer after three rounds of selection show significant enrichment in Round 2 with 1µM or 100 nM Caspase-3 treatment. The number of released phage was determined by counting infectious units as a function of round of selection.

**Supplementary Figure 6.**
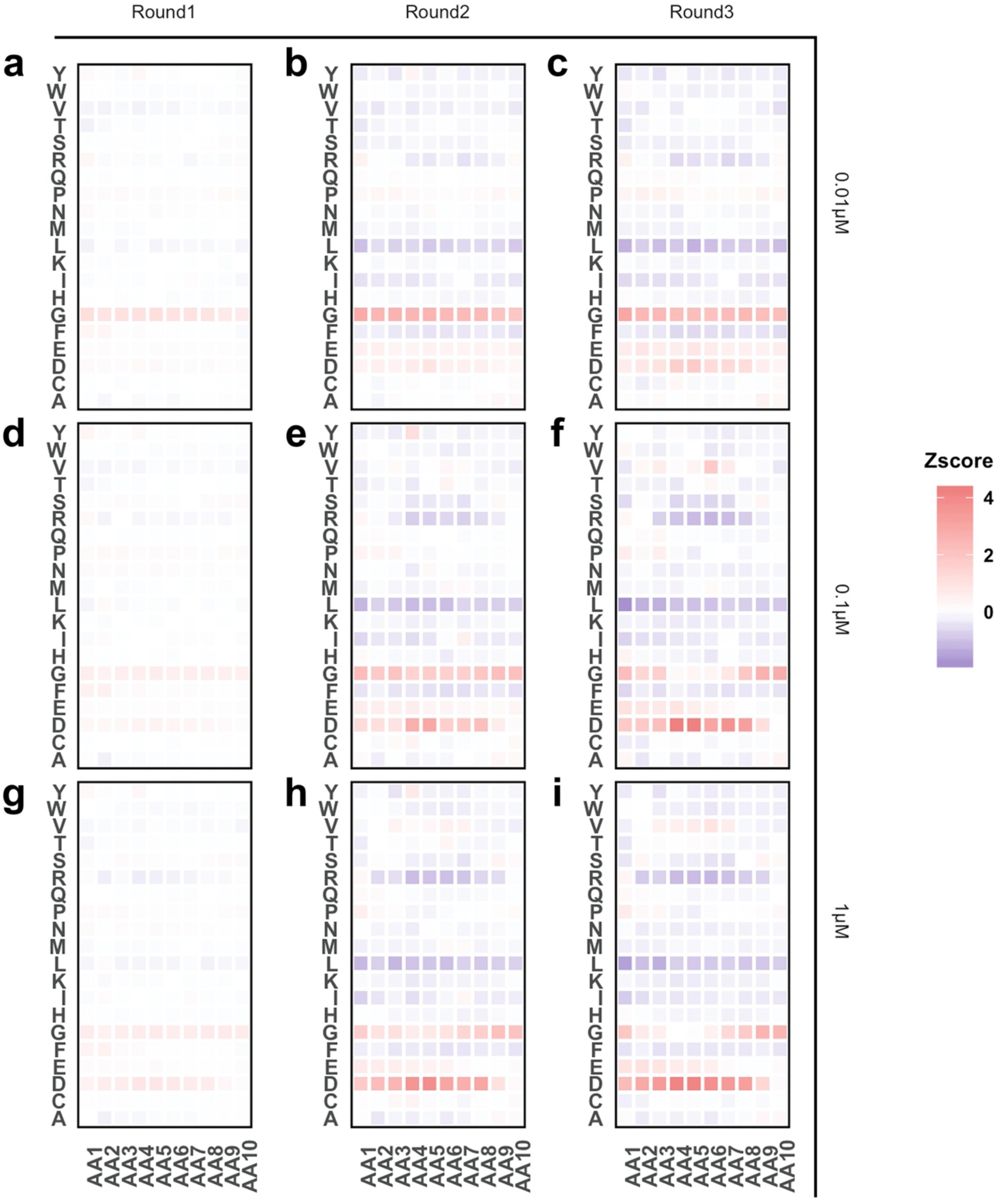
Heatmap reveals the amino acid composition change compared to the input library after three rounds of selection.

**Supplementary Figure 7.**
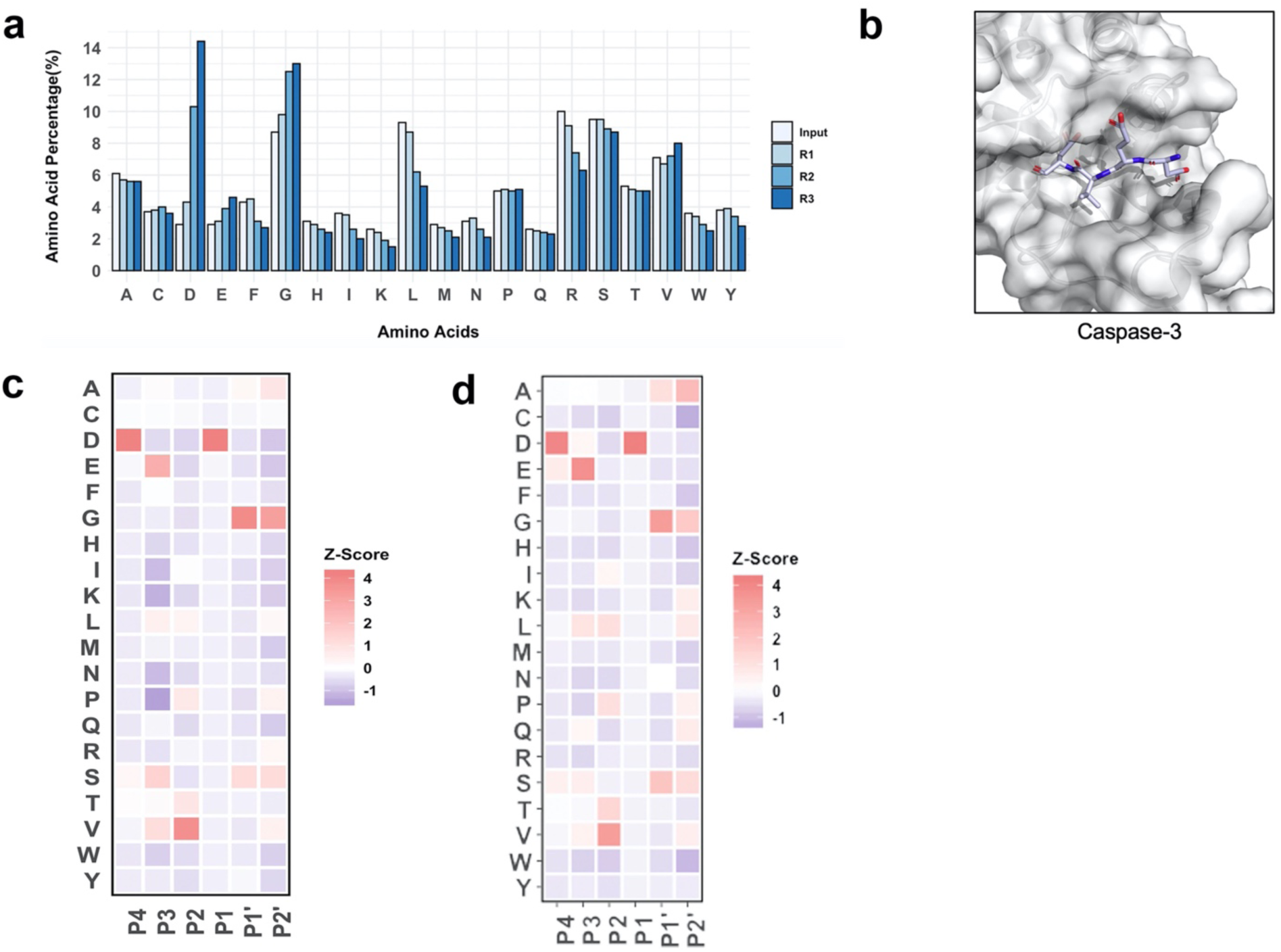
(a) Overall amino acid composition of the input library (Lib 10AA) and the outputs of Round 1, 2, 3. (b) Crystal structure of Caspase-3 with DEVD substrate binding to the pocket (adapted from PDB: 3PD1). (c, d) The heatmap for caspase-3 substrates generated based on top 20,000 sequences identified from screening Lib 10AA (Round 3, c) and from MEROPS dataset (d).

**Supplementary Figure 8.**
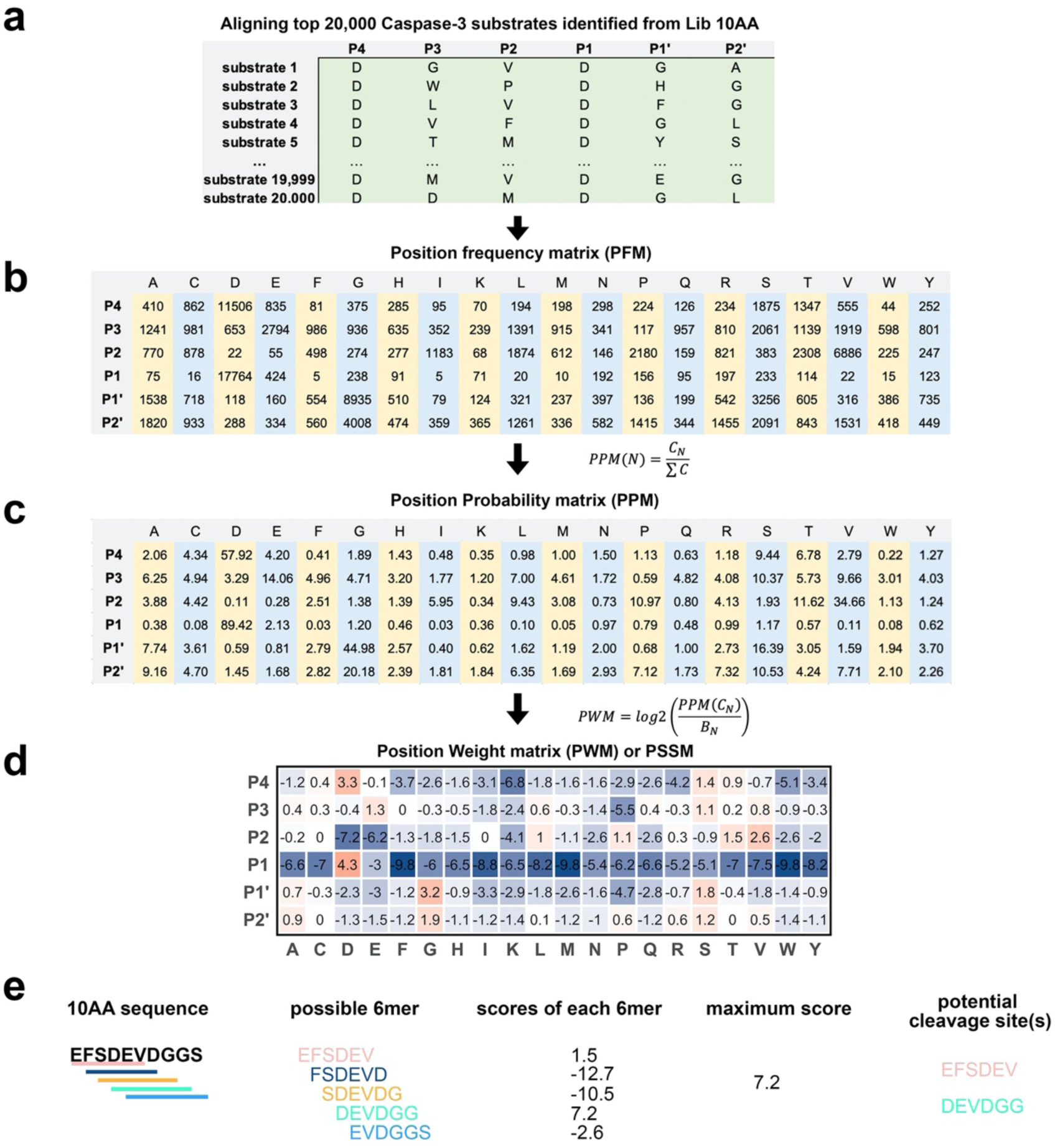
Generation of the PSSM. To illustrate the process, we use Caspase-3 as an example. (a) Top 20,000 potential Caspase-3 substrates identified from Lib 10AA (Round 3) were aligned. (b) To more accurately reflect the characteristics at each position, a matrix that contains the number of observed amino acids at each position is calculated, known as PFM. (c) A PPM was created based on PFM. (d) The PPM was converted to a PWM using a formula that converts it to a log-scale. PWMs are also known as position-specific scoring matrices (PSSM). (e) Identification of potential cleavage site(s).

**Supplementary Figure 9.**
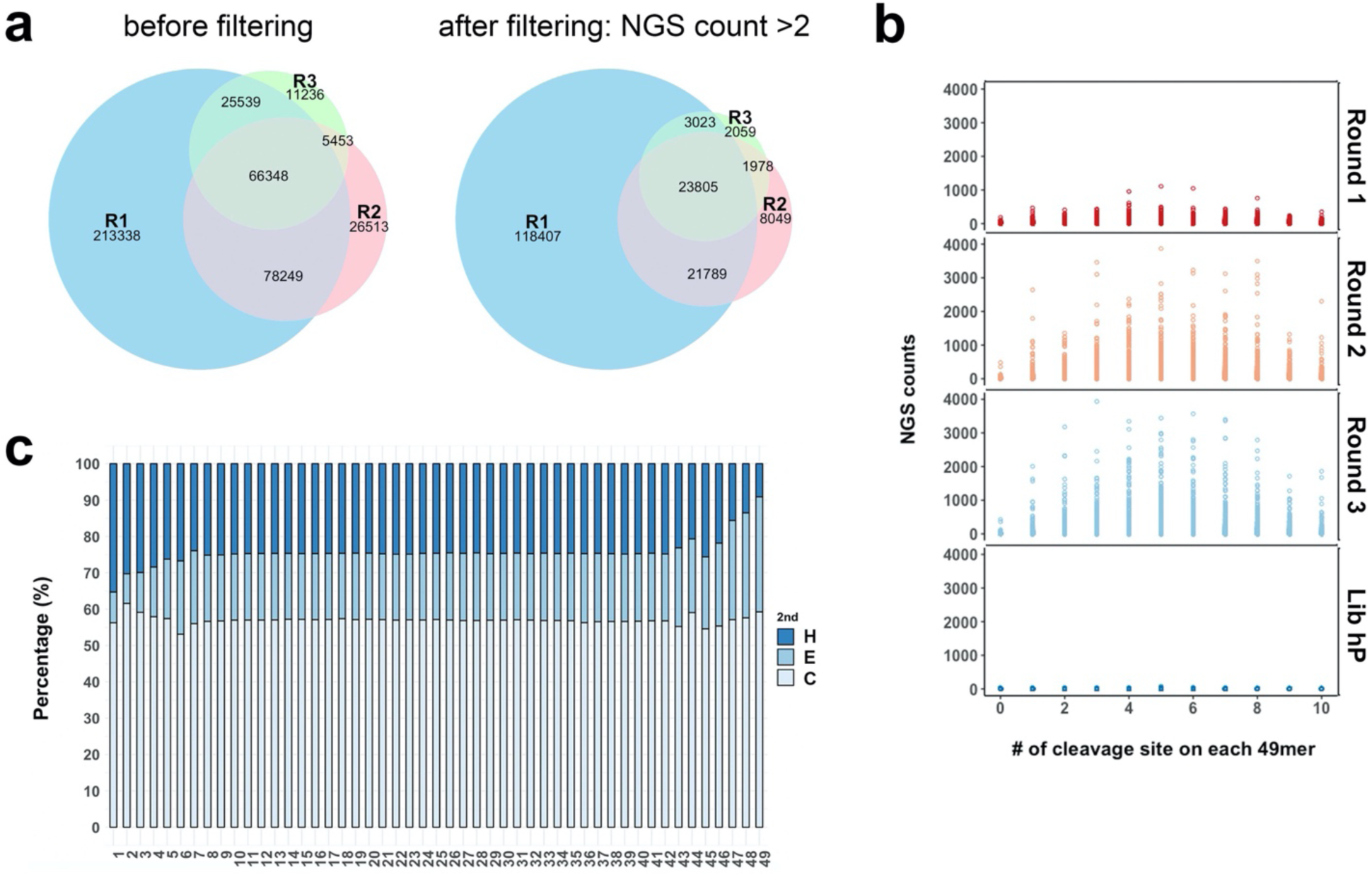
(a) The Venn diagram of unique peptides identified from Lib hP that decrease due to increasing enrichment with each of three rounds of the selection. Sequences with low counts (i.e., <4 counts) being pre-filtered out increases the overlap of the sequences in three rounds. (b) Roughly, the 49mer with more cleavage sites on it got enriched faster. NGS count was affected by many other factors, like PCR, phage amplification by E coli and whether the cleavage site is a good substrate. (c) The distribution of three different structures throughout the whole input library. The GOR (Garnier-Osguthorpe-Robson) method was used for 2nd structure prediction. The secondary structure throughout the library is comparably even but the calculation result of the C- and N-terminal of 49mer is affected by the constant flanking regions.

**Supplementary Figure 10.**
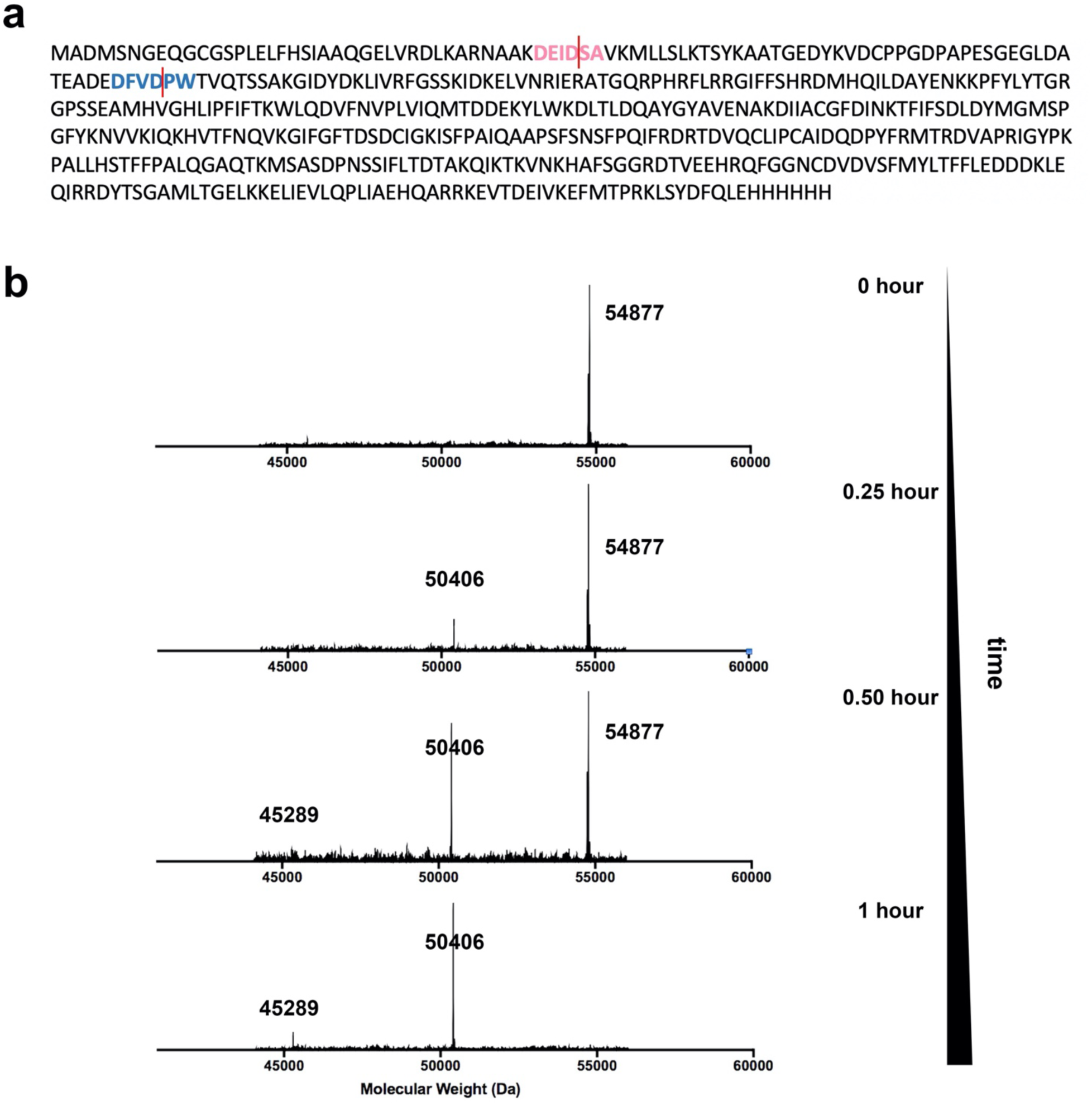
(a) The protein sequence of WARS. Two caspase-3 cleavage sites identified in substrate phage screening were highlighted in pink and red. (b) Representative ESI mass spectra of the intact protein and the fragments after caspase-3 proteolysis.

**Supplementary Figure 11.**
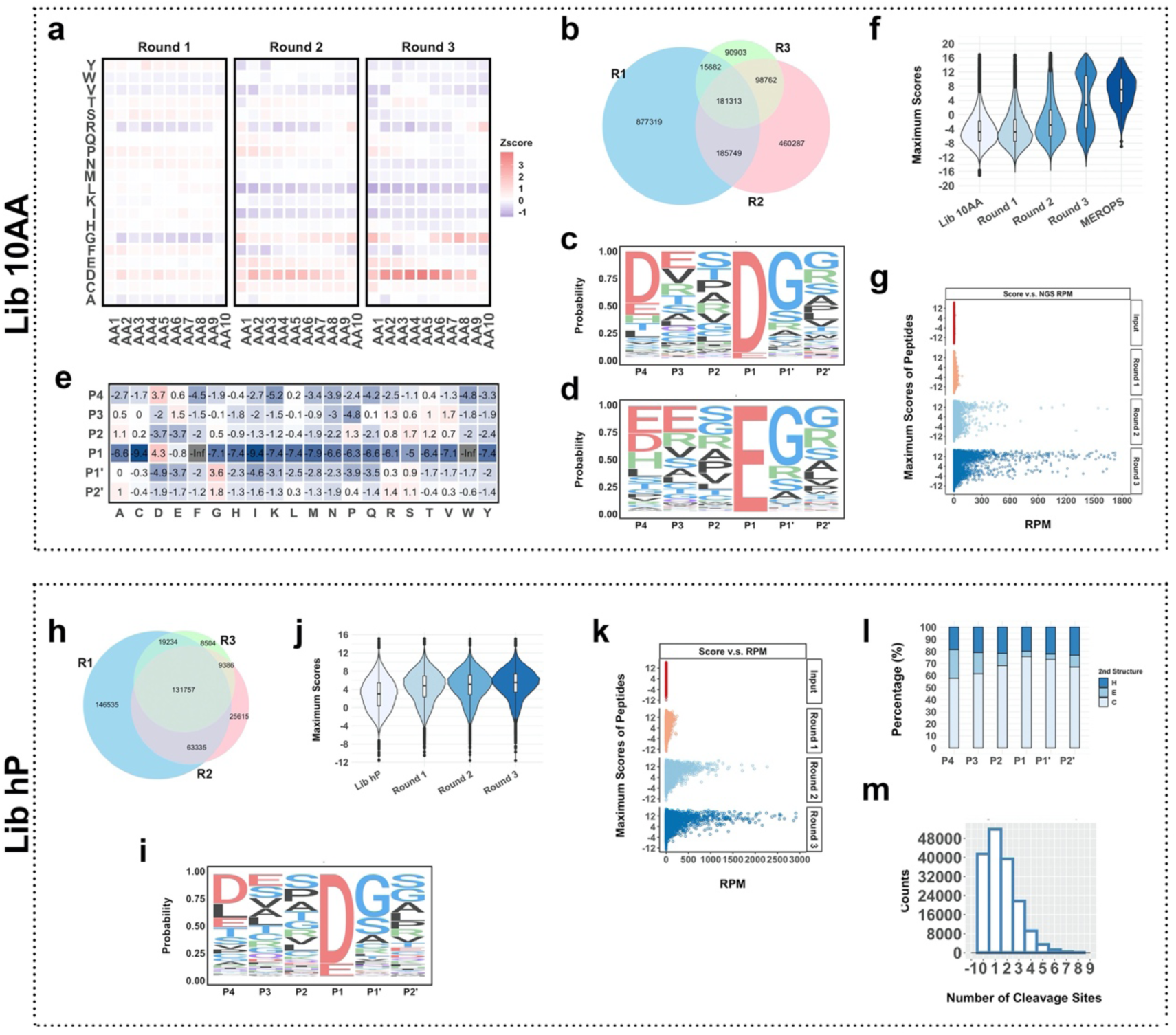
SPD-NGS libraries to identify protease specificity for Caspase-2. (a) Heatmap reveals the amino acid composition change compared to the input library after three rounds of selection. (b) The Venn diagram of unique peptides identified from Lib 10AA as a function of rounds of selection. (c) The Sequence Logo (Probability) for caspase substrates generated by aligning to top ∼20,000 sequences identified from screening Lib 10AA (Round 3). (d) A similar motif is obtained by selecting only proteolytic sites with a glutamate (E) at P1 position as seen previously by N-terminomics data of caspase-6. (e) The PSSM of caspase substrates is generated according to the alignment. (f) Violin plot of the maximum scores of 10AA peptides in input library, Round 1, 2, 3 outputs and the scores of peptides in MEROPS proteomics database. (g) The plot of maximum score vs NGS RPM for peptides in input library and Round 1, 2, 3 outputs. The peptides with higher scores get enriched faster. (h) The Venn diagram of unique peptides identified from Lib hP decreases with each of three rounds of the selection as enrichment increases as was seen for Lib 10AA. (i) The Sequence Logo (Probability) of caspase substrate consensus generated by all cleavage events identified from Lib hP. (j) Violin plot of the maximum scores of the Lib hP input library, and progressive enrichment of substrates as one progresses from output of Round 1, 2, and 3. (k) The plot of maximum score vs NGS RPM for peptides in input library and Round 1, 2, and 3 outputs. (l) Caspase has a known structural preference for cutting loop>helix>sheets and this matches structural bioinformatics for sites identified from Lib hP. P1method was used for 2nd structure prediction. (m) Distribution of frequency observed as a function of number of cuts in the 49AA peptides show most have one cut but some have multiple that decays monotonically.

**Supplementary Figure 12.**
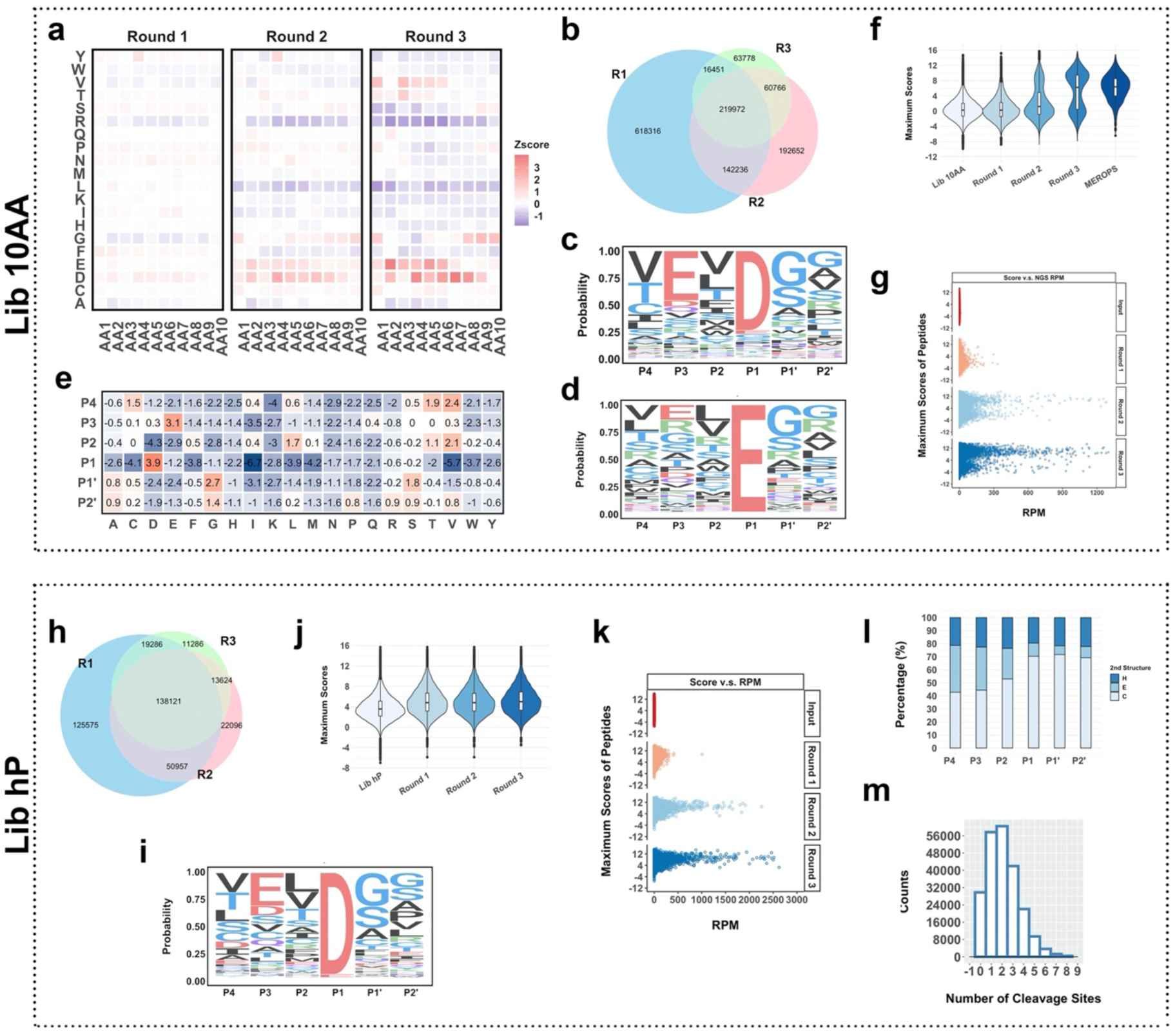
SPD-NGS libraries to identify protease specificity for Caspase-6. (a) Heatmap reveals the amino acid composition change compared to the input library after three rounds of selection. (b) The Venn diagram of unique peptides identified from Lib 10AA as a function of rounds of selection. (c) The Sequence Logo (Probability) for caspase substrates generated by aligning to top ∼20,000 sequences identified from screening Lib 10AA (Round 3). (d) A similar motif is obtained by selecting only proteolytic sites with a glutamate (E) at P1 position as seen previously by N-terminomics data of caspase-6. (e) The PSSM of caspase substrates is generated according to the alignment. (f) Violin plot of the maximum scores of 10AA peptides in input library, Round 1, 2, 3 outputs and the scores of peptides in MEROPS proteomics database. (g) The plot of maximum score vs NGS RPM for peptides in input library and Round 1, 2, 3 outputs. The peptides with higher scores get enriched faster. (h) The Venn diagram of unique peptides identified from Lib hP decreases with each of three rounds of the selection as enrichment increases as was seen for Lib 10AA. (i) The Sequence Logo (Probability) of caspase substrate consensus generated by all cleavage events identified from Lib hP. (j) Violin plot of the maximum scores of the Lib hP input library, and progressive enrichment of substrates as one progresses from output of Round 1, 2, and 3. (k) The plot of maximum score vs NGS RPM for peptides in input library and Round 1, 2, and 3 outputs. (l) Caspase has a known structural preference for cutting loop>helix>sheets and this matches structural bioinformatics for sites identified from Lib hP. P1method was used for 2nd structure prediction. (m) Distribution of frequency observed as a function of number of cuts in the 49AA peptides show most have one cut but some have multiple that decays monotonically.

**Supplementary Figure 13.**
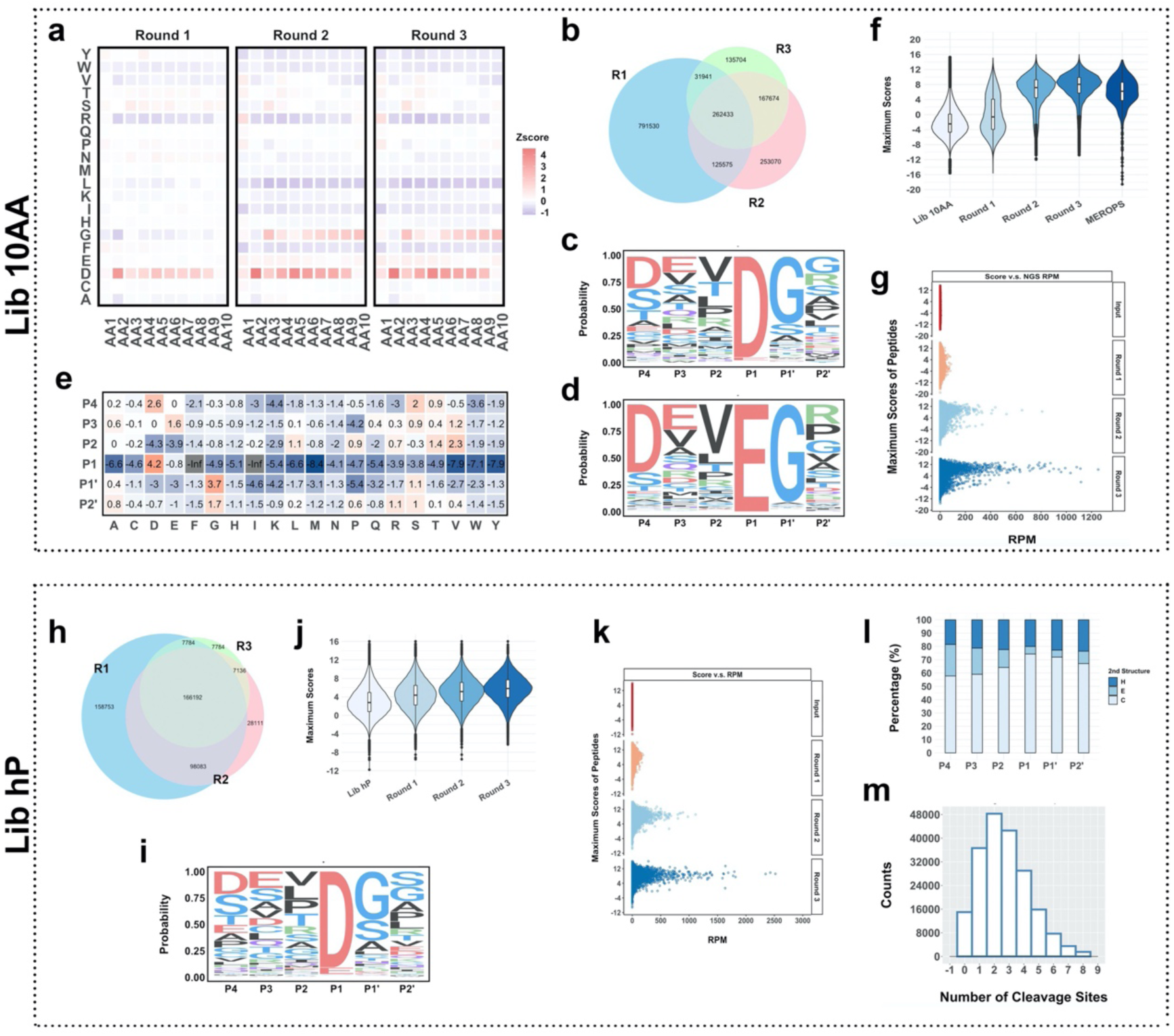
SPD-NGS libraries to identify protease specificity for Caspase-7. (a) Heatmap reveals the amino acid composition change compared to the input library after three rounds of selection. (b) The Venn diagram of unique peptides identified from Lib 10AA as a function of rounds of selection. (c) The Sequence Logo (Probability) for caspase substrates generated by aligning to top ∼20,000 sequences identified from screening Lib 10AA (Round 3). (d) A similar motif is obtained by selecting only proteolytic sites with a glutamate (E) at P1 position as seen previously by N-terminomics data of caspase-6. (e) The PSSM of caspase substrates is generated according to the alignment. (f) Violin plot of the maximum scores of 10AA peptides in input library, Round 1, 2, 3 outputs and the scores of peptides in MEROPS proteomics database. (g) The plot of maximum score vs NGS RPM for peptides in input library and Round 1, 2, 3 outputs. The peptides with higher scores get enriched faster. (h) The Venn diagram of unique peptides identified from Lib hP decreases with each of three rounds of the selection as enrichment increases as was seen for Lib 10AA. (i) The Sequence Logo (Probability) of caspase substrate consensus generated by all cleavage events identified from Lib hP. (j) Violin plot of the maximum scores of the Lib hP input library, and progressive enrichment of substrates as one progresses from output of Round 1, 2, and 3. (k) The plot of maximum score vs NGS RPM for peptides in input library and Round 1, 2, and 3 outputs. (l) Caspase has a known structural preference for cutting loop>helix>sheets and this matches structural bioinformatics for sites identified from Lib hP. P1method was used for 2nd structure prediction. (m) Distribution of frequency observed as a function of number of cuts in the 49AA peptides show most have one cut but some have multiple that decays monotonically.

**Supplementary Figure 14.**
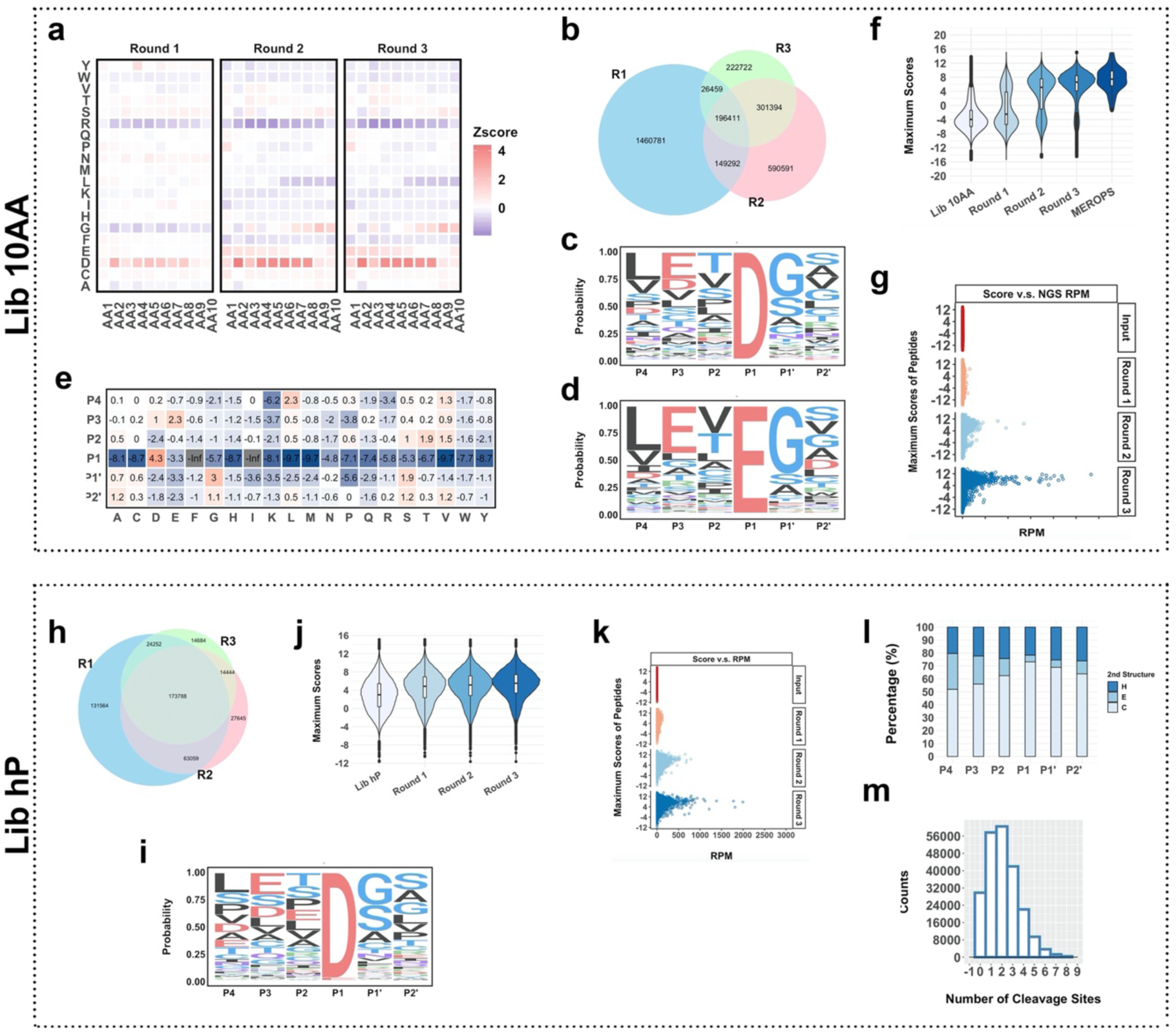
SPD-NGS libraries to identify protease specificity for Caspase-8. (a) Heatmap reveals the amino acid composition change compared to the input library after three rounds of selection. (b) The Venn diagram of unique peptides identified from Lib 10AA as a function of rounds of selection. (c) The Sequence Logo (Probability) for caspase substrates generated by aligning to top ∼20,000 sequences identified from screening Lib 10AA (Round 3). (d) A similar motif is obtained by selecting only proteolytic sites with a glutamate (E) at P1 position as seen previously by N-terminomics data of caspase-6. (e) The PSSM of caspase substrates is generated according to the alignment. (f) Violin plot of the maximum scores of 10AA peptides in input library, Round 1, 2, 3 outputs and the scores of peptides in MEROPS proteomics database. (g) The plot of maximum score vs NGS RPM for peptides in input library and Round 1, 2, 3 outputs. The peptides with higher scores get enriched faster. (h) The Venn diagram of unique peptides identified from Lib hP decreases with each of three rounds of the selection as enrichment increases as was seen for Lib 10AA. (i) The Sequence Logo (Probability) of caspase substrate consensus generated by all cleavage events identified from Lib hP. (j) Violin plot of the maximum scores of the Lib hP input library, and progressive enrichment of substrates as one progresses from output of Round 1, 2, and 3. (k) The plot of maximum score vs NGS RPM for peptides in input library and Round 1, 2, and 3 outputs. (l) Caspase has a known structural preference for cutting loop>helix>sheets and this matches structural bioinformatics for sites identified from Lib hP. P1method was used for 2nd structure prediction. (m) Distribution of frequency observed as a function of number of cuts in the 49AA peptides show most have one cut but some have multiple that decays monotonically.

**Supplementary Figure 15.**
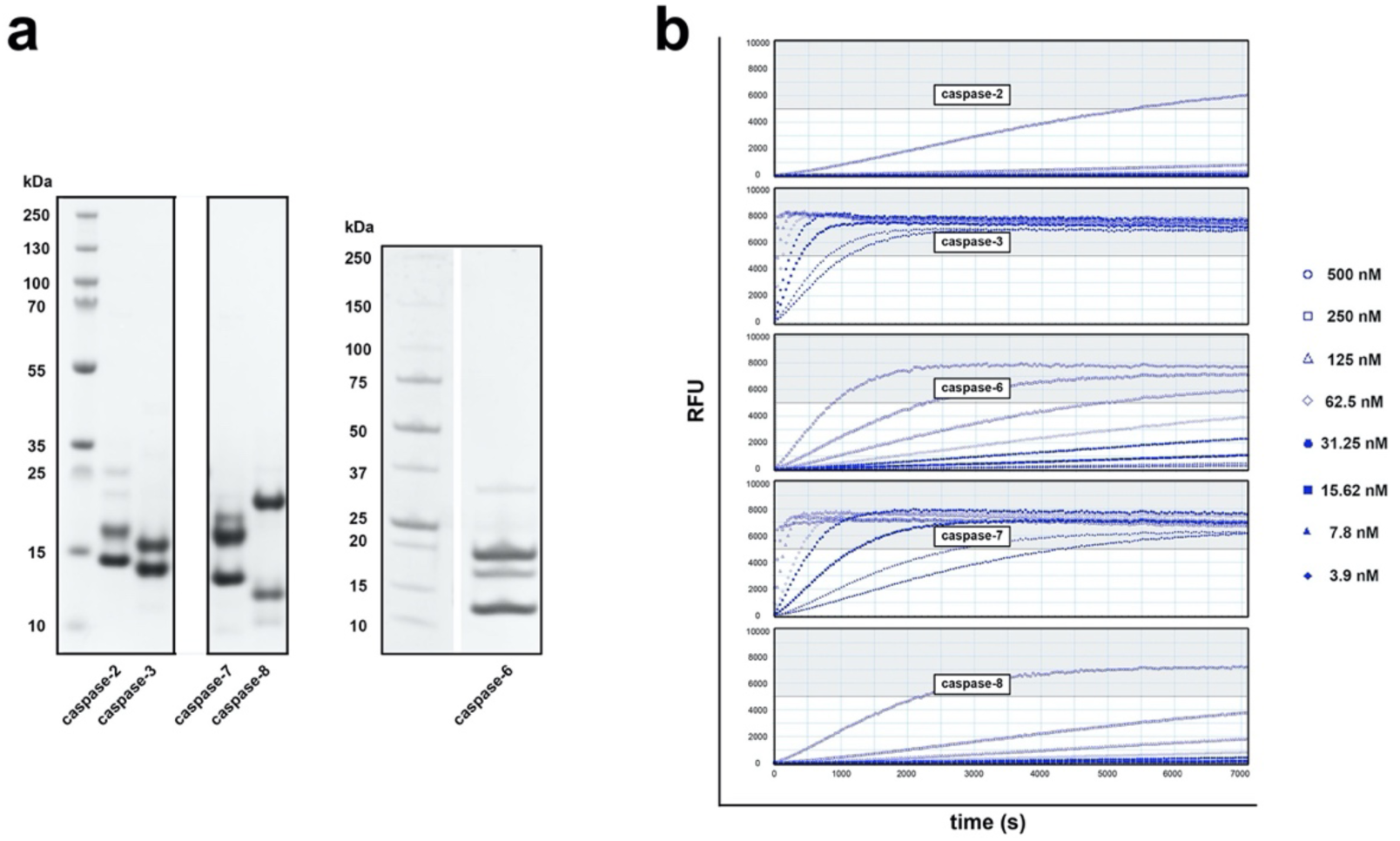
(a) Gel electrophoresis of active recombinant caspase-2, -3, -6, -7 and -8 purified from E. coli. b. (b) The activity of purified active recombinant caspases-2, -3, -6, -7 and -8 in caspase activity buffer. Ac-DEVD-R110 was used to monitor the caspase activity.

**Supplementary Figure 16.**
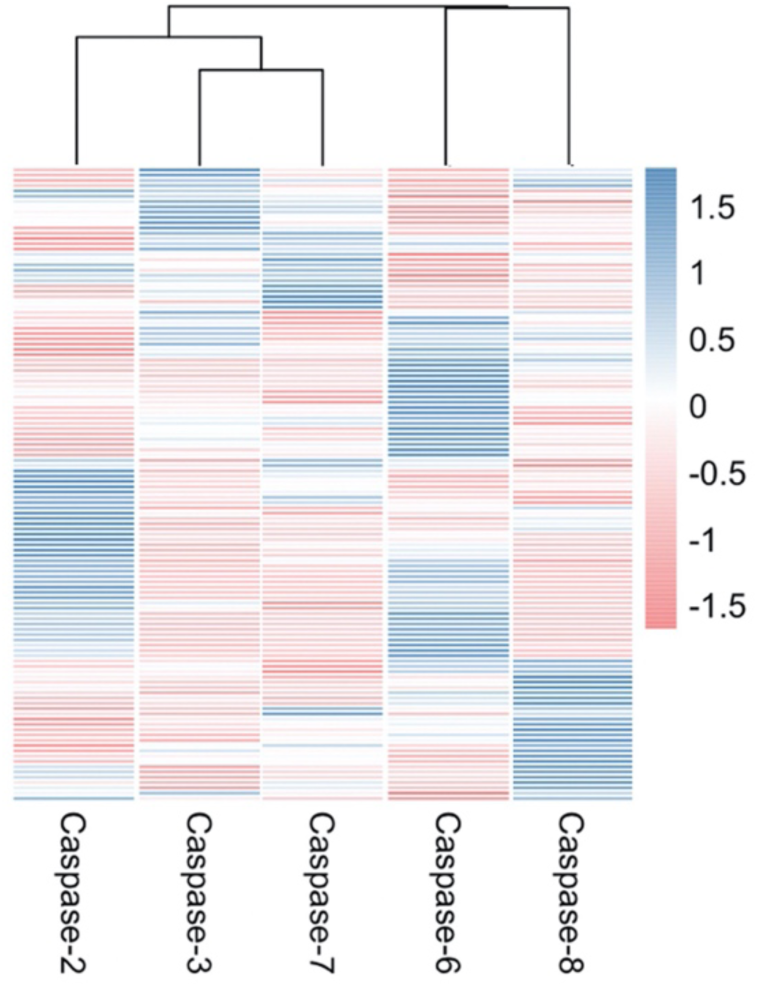
Cluster the heat maps of each

**Supplementary Figure 17.**
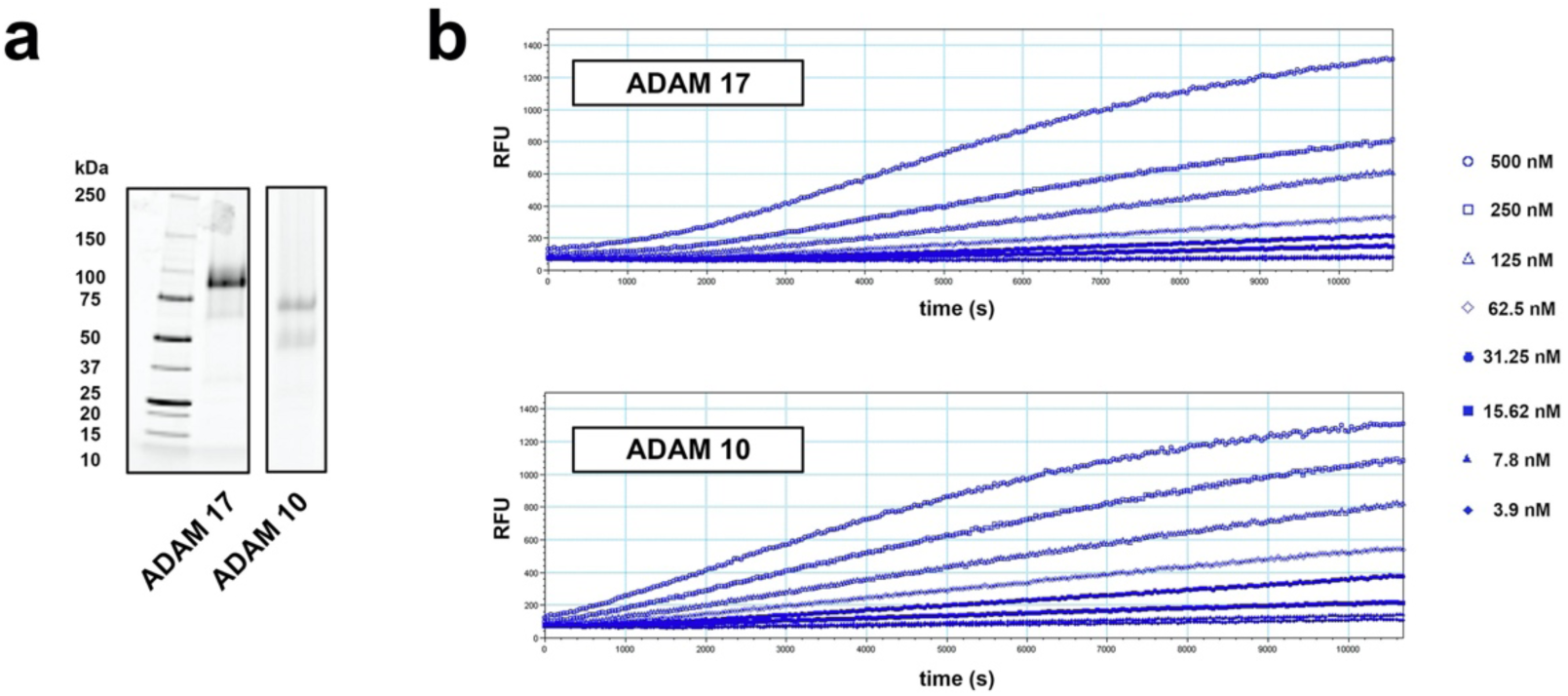
(a) Gel electrophoresis of active recombinant active ADAM 10 and ADAM 17 expressed by Expi293 cells. (b) The activity of purified active recombinant ADAM10 and ADAM 17 in activity buffer. Mca-KPLGL-Dpa-AR-NH2 was used to monitor the ADAMs activity.

**Supplementary Figure 18.**
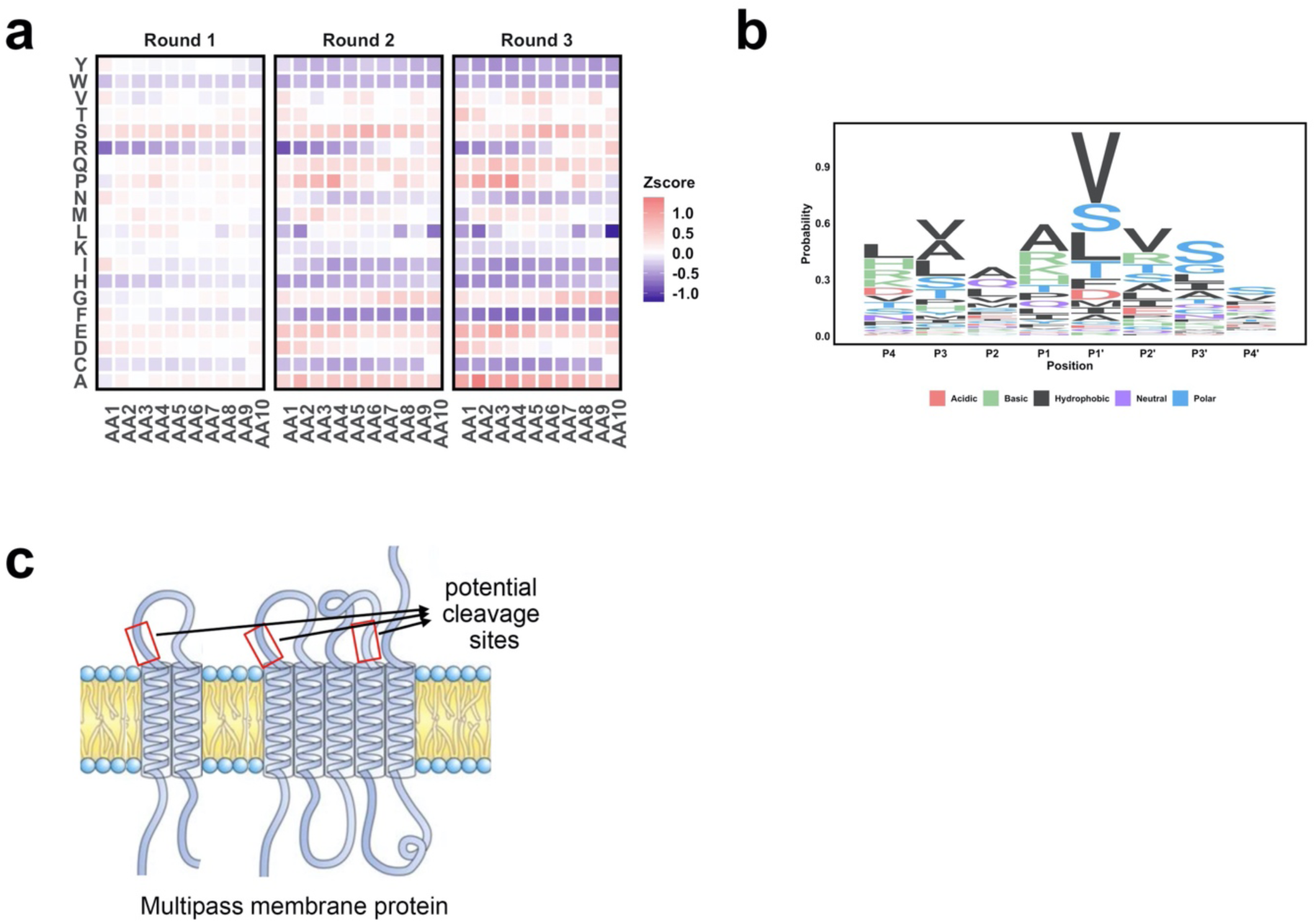
SPD-NGS libraries to identify protease specificity for ADAM 17. (a) Heatmap reveals the amino acid composition change compared to the input library after three rounds of selection. (b) The Sequence Logo (Bits) for ADAM17 substrates generated by substrate sequences recorded in MEROPS database.

**Supplementary Figure 19.**
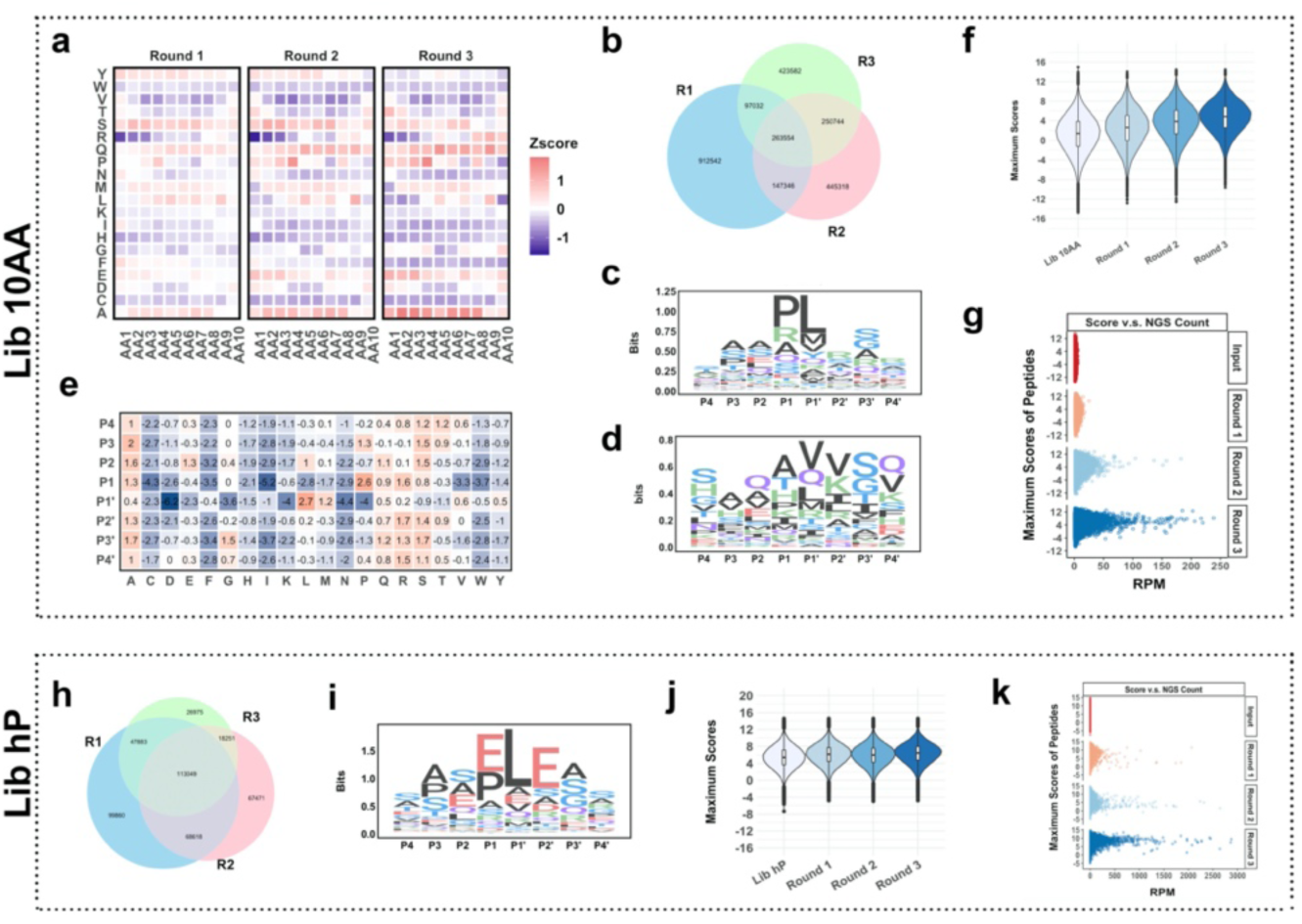
SPD-NGS libraries to identify protease specificity for ADAM 10. (a) Heatmap reveals the amino acid composition change compared to the input library after three rounds of selection. (b) The Venn diagram of unique peptides identified from Lib 10AA as a function of rounds of selection. (c) The Sequence Logo (Bits) for ADAM10 substrates generated by aligning the top ∼20,000 sequences identified from screening Lib 10AA (Round 3). (d) The Sequence Logo (Bits) for ADAM10 substrates recorded in MEROPS database. (e) The PSSM of ADAM10 substrates is generated according to the alignment. (f) Violin plot of the maximum scores of 10AA peptides in input library, Round 1, 2, 3 outputs and the scores of peptides in MEROPS proteomics database. (g) The plot of maximum score vs NGS RPM for peptides in input library and Round 1, 2, 3 outputs. The peptides with higher scores get enriched faster. (h) The Venn diagram of unique peptides identified from Lib hP decreases with each of three rounds of the selection as enrichment increases as was seen for Lib 10AA. (i) The Sequence Logo (Bits) of ADAM10 substrate consensus generated by all cleavage events identified from Lib hP. (j) Violin plot of the maximum scores of the Lib hP input library, and progressive enrichment of substrates as one progresses from output of Round 1, 2, and 3. (k) The plot of maximum score vs NGS RPM for peptides in input library and Round 1, 2, and 3 outputs.

**Supplementary Figure 20.**
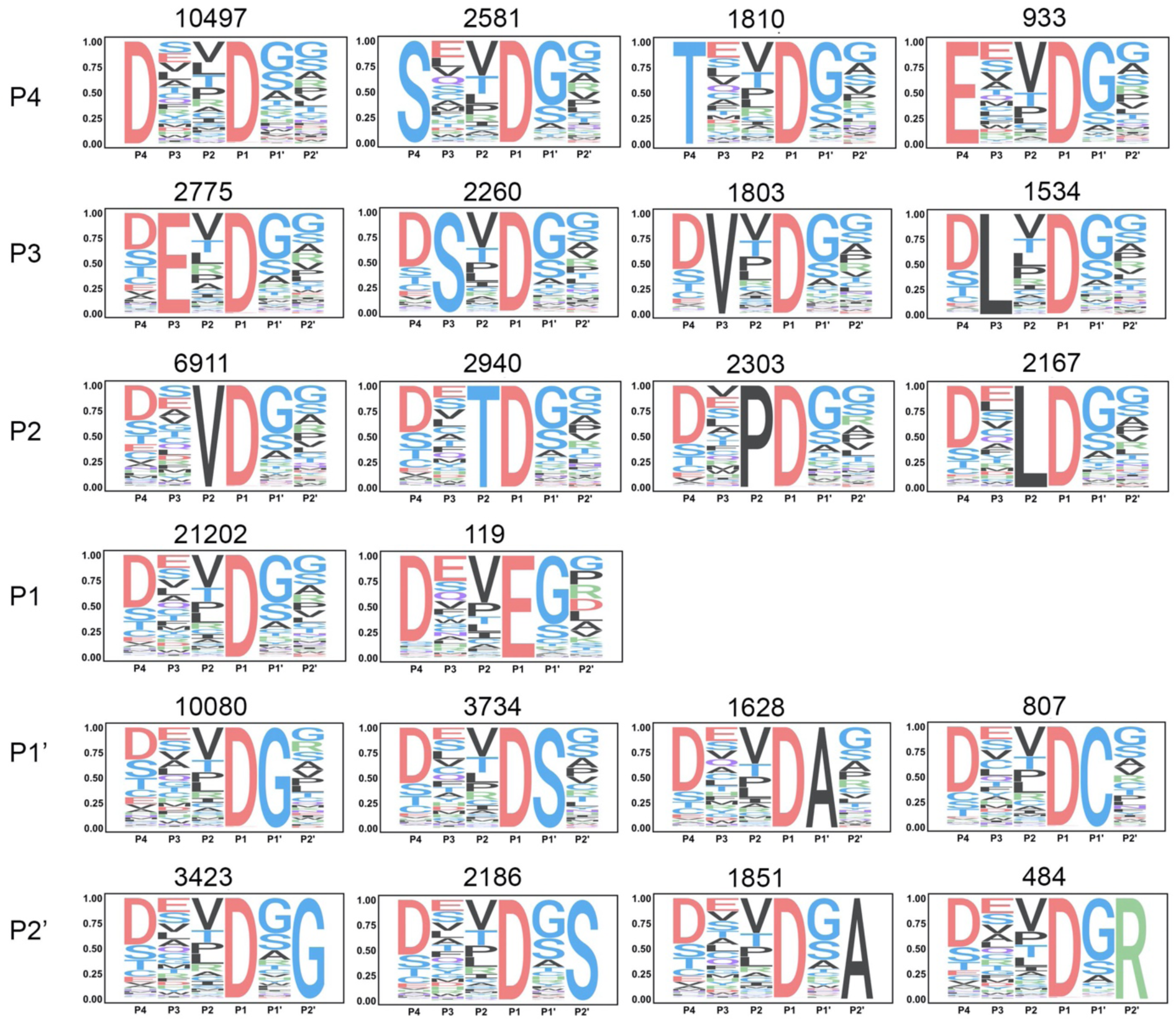
The Sequence Logo (Probability) for caspase substrates generated with the alignment of top ∼ 20,000 sequences enriched from SPD-NGS with Lib 10AA. Each of the sequence logo is generated by fixing the top abundant residues at different positions. The number above the sequence logo indicates the number of the sequences used for generating the logo.

**Supplementary Figure 21.**
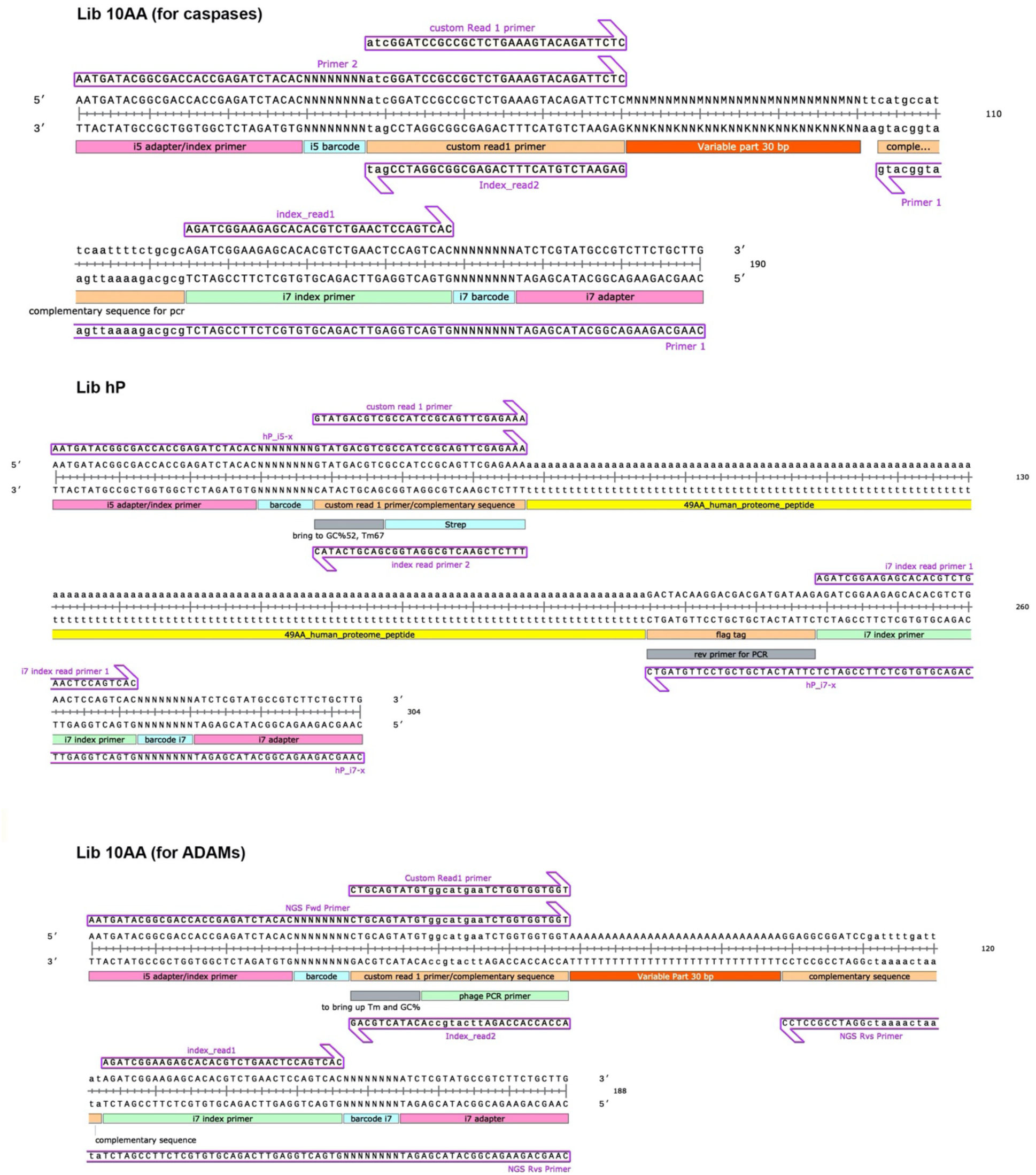
Primer design for NGS

